# Human neurons undergo protracted functional maturation into adulthood

**DOI:** 10.1101/2025.08.01.668139

**Authors:** Matthijs B. Verhoog, Roxanne M. Hattingh, Pia Scharein, Tariro A. Chatiza, Nawaal Samsodien, Yanelisa Pulani, Amalia N. Awala, Jared Tavares, Eline J. Mertens, Christiaan P. J. De Kock, Björn Kampa, Tim Kroon, Huibert D. Mansvelder, Ursula K. Rohlwink, Anthony Figaji, Graham Fieggen, Sean Tromp, Johannes M. N. Enslin, Roger Melvill, James Butler, Joseph V. Raimondo

## Abstract

Human cognitive development is uniquely prolonged^1–3^, reflecting the extended postnatal maturation of the cerebral cortex where cell-type differentiation^4,5^, synaptogenesis^6,7^, myelination^8^ and transcriptional regulation^5,9,10^ all follow protracted developmental timelines. However, when the intrinsic physiological properties of human cortical neurons mature, and how this compares with other species, remains unknown. Here we show, through patch-clamp recordings of temporal cortex from an ancestrally diverse cohort spanning infancy to adulthood, that human supragranular pyramidal neurons exhibit pronounced neoteny of their functional properties, maturing well into adult life. In the mouse the same sequence of changes unfolds within weeks and scales tightly with brain volume. In humans this allometric relationship is fundamentally different, with the trajectories of individual properties continuing for decades beyond peak brain volume, up to 7 times more slowly than the difference in brain growth rates alone would predict.

Together, these trajectories define a continuous developmental axis along which the neuronal population progresses throughout cognitive development, with the neurons present at each stage differing in how they transform their inputs. Notably, the population diverges only in adolescence, when one of its two adult end-states acquires a distinctive physiology, absent in mice, that characterises the deep supragranular neurons linking distant cortical areas in humans^11,12^ and other primates^13–15^. A generative model of the developing human supragranular population, unifying the developmental models of every physiological domain, now captures this protracted maturation from infancy to adulthood, providing a resource for simulation, normative referencing and cross-species age mapping.

---

Across mammals, larger brains and the greater cognitive capacities they support are generally accompanied by longer periods of postnatal development, and in this the human brain is an extreme outlier^1,2^. What unfolds over this protracted period has been charted anatomically^7,8^ and molecularly^4–6,9,10^, but how neuronal intrinsic physiological properties mature in the intact human brain, and over what time course, has remained largely uncharacterized. We therefore set out to define this trajectory, to relate it to the volumetric growth of the brain, and to compare it with the mouse. As the principal model organism of cortical neuroscience, and a species that reaches maturity within weeks, the mouse sits at the opposite extreme of this developmental spectrum, which makes it an especially informative comparator when establishing how far the neoteny of human neurons extends into their functional physiology.

## Results

### Human neurons mature over decades

To establish the nature and time course of physiological maturation in human cortical neurons, we performed whole-cell patch-clamp recordings in surgically resected brain tissue from subjects spanning infancy to adulthood (n = 584 cells, 80 subjects, 10 months to 55 years, Fig. 1a, Fig. S1, Table S1-2). We focused exclusively on pyramidal neurons in the upper, ‘supragranular’ layers of temporal association cortex, which exhibit enriched glutamatergic cell-type diversity with emerging human-specific specialisations and developmentally regulated programs linked to cognition^12,16,17^. Using generalized additive mixed models (GAMMs) to capture the developmental trajectory of each electrophysiological feature (Fig. 1b, Fig. S2-5), we found that the physiology of human supragranular neurons continues to mature well into adulthood (Fig. 2a, Fig. S7a).

**Figure 1.**
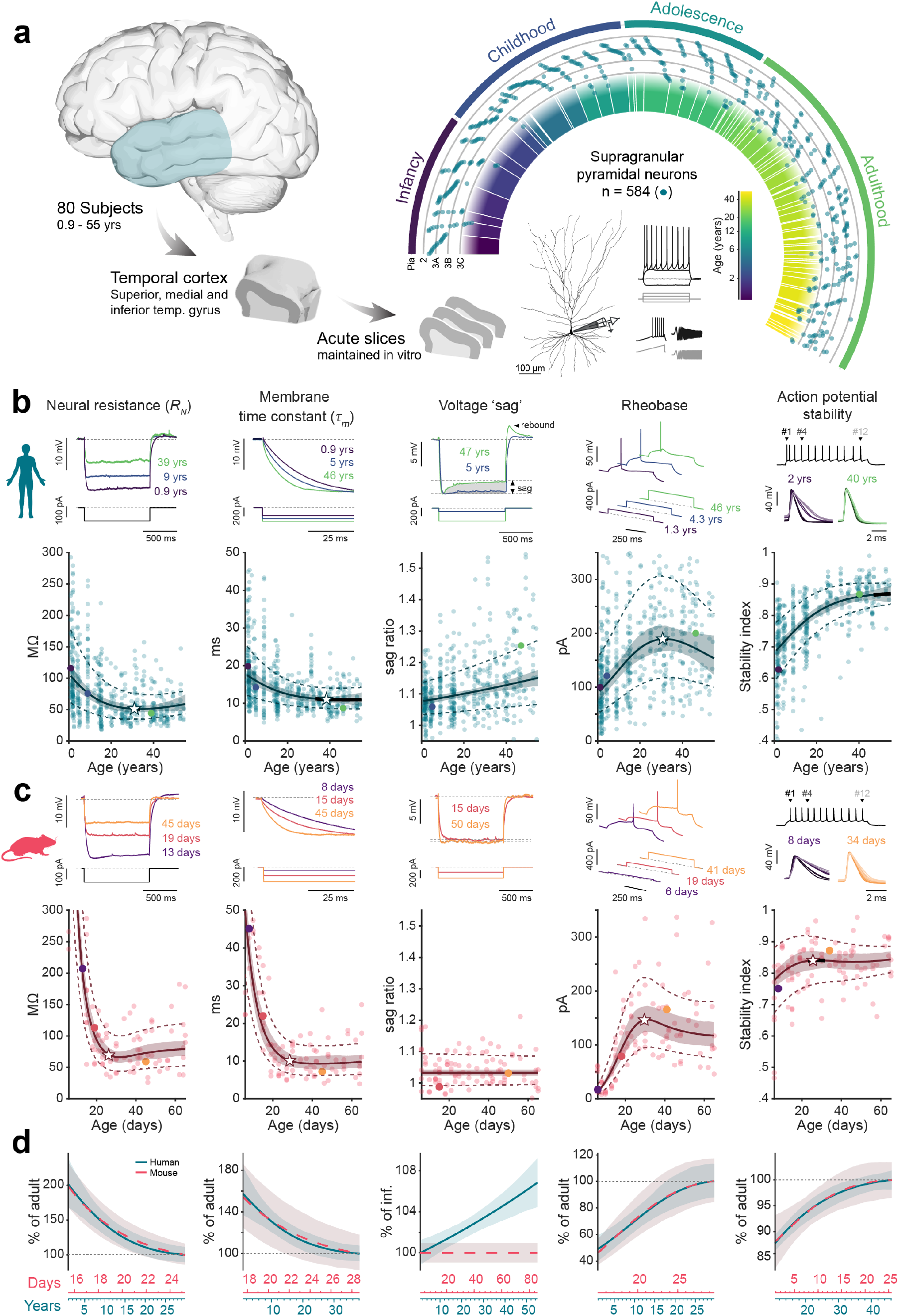
Human cortical pyramidal neurons have extended postnatal maturation trajectories. **a**, Experimental pipeline for characterization of human pyramidal neuron physiology in neurosurgical samples of temporal cortex. Neurons were targeted for whole-cell patch-clamp recordings throughout supragranular layers 2-3C, from infancy to adulthood. Arc shows distribution of 584 recorded neurons across ages and cortical depths (radius, 0-1500 μm). **b**, Developmental trajectories of human neural properties fit with GAMMs, each point represents measure from a single neuron. Predicted mean and 95% confidence interval (c.i.) shown by solid line and shading; dashed lines show the fitted standard deviation. Star and thick line indicate the maturation peak age and its IQR (Fig. 2a). Top traces show representative examples at different ages, with the corresponding data points in matching colours below. Some y-axes are truncated for display, full datasets presented in Fig. S2-S4. **c**, As **b**, for L2/3 pyramidal neurons in mouse temporal association cortex. **d**, Human and mouse trajectories overlaid, each expressed as a percentage of its own adult value (+/-95% c.i.). The two age axes are scaled so that the curves superimpose, aligning ages equally far from adult value in both species. Sag ratio does not change with age in mouse and is shown as percentage of its infant value

**Figure 2.**
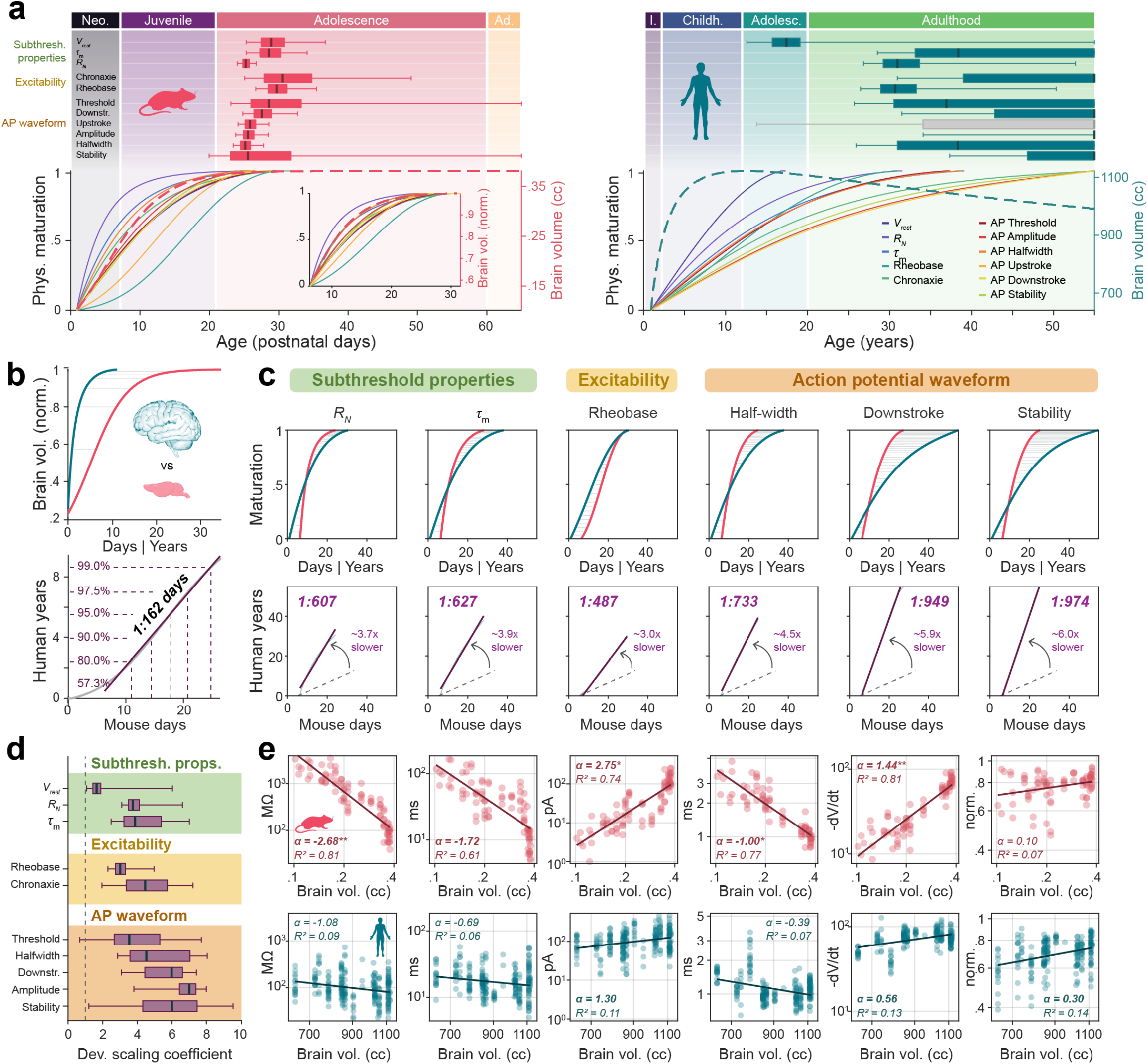
Human physiological development is desynchronised from brain growth. **a**, Top: maturation peak ages of mouse (left) and human (right) physiological features, grouped by physiological domain. Boxes show the bootstrap median and IQR, whiskers the 2.5th and 97.5th percentiles; grey, no significant age effect. Bottom: normalized developmental trajectories of the same features, with brain volume growth (dashed) on the same age axis; shaded bands mark developmental stages. Inset: mouse trajectories over same proportional brain growth range sampled in humans. **b**, Dynamic time warping (DTW) of human and mouse brain growth curves, grey lines show examples of matched points. Below: warping path (grey) with linear fit to yield brain growth scale factor; dashed lines link ages at indicated % of adult brain volume. **c**, As **b**, for physiological features, with scale factor given in days. Dashed grey line shows the brain growth slope (**b**), along with the developmental scaling coefficient. **d**, Developmental scaling coefficients for all features with a significant age effect in both species (bootstrap median, IQR and 2.5th to 97.5th percentiles). **e**, Allometric relationships between physiological features and brain volume in mouse (top) and human (bottom) over the brain growth phase; allometric scaling exponent α is slope of LMM fit on log-log axes (Table S13), shown with its marginal *R²*; bold α, 95% c.i. excludes zero; asterisks, significant species difference (\**P* < 0.05, *\*\*P < 0.01*, \*\*\**P* < 0.001).

These maturational changes extended to every physiological domain, from how a neuron integrates its inputs to its excitability and action potential firing. Membrane resistance and time constant, which set a neuron’s sensitivity to inputs and its temporal integration, decreased slowly and steadily with age, halving over the first three decades of life (Fig. 1b, Fig. S2). The GAMMs model the variance alongside the mean, which revealed a pronounced developmental decrease in variability (Fig. 1b, Fig. S2f, Table S3-S4), primarily reflecting changes in within-subject variance (Fig. S2f-h).

Adult human supragranular neurons express HCN channels, which filter slow subthreshold inputs and produce the characteristic voltage ‘sag’ on hyperpolarization^11,18^. This feature was minimally present during childhood and progressively increased through adulthood (Fig. 1b). Notably, as sag ratio increased, so did its variability, suggesting late HCN channel expression contributes to adult physiological diversity (Fig. 1b, Fig. S2f,g). In line with a growing influence of HCN channels on neuronal membrane properties, both resonant frequency and cutoff frequency increased significantly with age (Fig. S12a,b), indicating that as they mature neurons become tuned to higher frequency oscillations and can process signals across a broader frequency range.

Excitability, too, changed gradually. Rheobase, the minimum current needed to elicit an action potential (AP), increased slowly over the first few decades of human life (Fig. 1b, Fig. S3) and the action potentials themselves changed as well. In juvenile neurons their waveform showed pronounced use-dependent dampening during repetitive firing, which resolved only towards middle adulthood (Fig. 1b, Fig. S4). This protracted timeline is particularly significant given that stable, rapid action potential kinetics have previously been linked to enhanced information processing and cognitive performance in humans^19–21^.

Throughout this progression, human neurons maintained their characteristic physiological heterogeneity^11,12,22–24^. Laminar position within the supragranular layers contributed substantially to this, with properties such as membrane sag varying systematically from superficial to deep^11,18^, and soma depth was a significant covariate in the GAMMs of most features (Table S3-10, Fig. S2e, Fig. S3d). Biological sex, evaluated in the same way, revealed no difference in the trajectories of male and female neurons (Fig. S5). The trajectories proved robust in a series of sensitivity analyses, remaining stable under leave-one-subject-out refitting, with unbiased cross-validated predictions across the age range (Fig. S6, Table S11), and under different model specifications (Fig. S9, Table S12).

To place this human timeline in a cross-species context, we recorded supragranular pyramidal neurons of mouse temporal association cortex across postnatal development and modelled them with the same approach (n = 151 cells, 42 mice, 0.9-86 days). With the exception of sag ratio, which remained absent in mice, the nature of physiological changes was clearly conserved: membrane resistance and time constant decreased, rheobase increased and action potentials stabilised in the same direction as in humans. These changes were again accompanied by developmental changes in variability (Fig. 1c, Fig. S2-S4), and most features reached mature levels within 3 to 4 weeks (Fig. 2a, Fig. S7a).

Together, these results map the developmental timeline of each feature in both species, revealing the same changes occur on substantially different timescales. Expressed relative to its own adult value, each feature’s trajectory superimposes across species (Fig. 1d, Fig. S4h). This gives a feature-by-feature correspondence of developmental age between mouse and human, defined by neuronal physiology itself. On this scale, weeks of mouse development correspond to years to decades of human life.

### Species-specific allometric rules

Larger brains develop for longer, so protracted human physiological maturation might be expected from our encephalization alone. Brain growth itself therefore provides the reference against which this timing can be judged. We compared the normalized developmental trajectories of physiological features with brain growth curves in both species. In mice, physiological maturation tracked brain growth closely, with most features reaching adult levels around the time maximum brain volume was attained, whereas in humans, maturation continued well beyond (Fig. 2a).

To quantify the relative maturation rates of the two species directly, we aligned the human and mouse developmental trajectories for brain volume and physiological features by dynamic time warping (DTW, Fig. 2b,c, Fig. S7b,c). The warping paths were consistently linear, so that the trajectories of the two species map onto each other through a constant scale factor. This factor varied by feature, for example one mouse day corresponded to 487 human days (1.3 years) for rheobase and to 733 human days (2 years) for AP half-width (Fig. 2c). Applied to the brain volume growth curves themselves, the same analysis gave a substantially smaller factor of 162 human days (0.4 years) per mouse day (Fig. 2b). Normalizing the physiological scale factors by this anatomical one showed that human neurophysiological maturation proceeds up to 7 times more slowly than brain growth alone would predict (Fig. 2d).

This differential scaling suggests a desynchronization of physiological and anatomical maturation in humans, possibly reflecting different underlying allometric growth rules. During development, an organism’s traits typically mature at predictable rates relative to one another, following power law relationships^25,26^. To test whether such relationships may exist between neural physiology and overall brain maturation, and whether they differ between species, we analyzed how physiological features vary with brain volume on logarithmic scales. Indeed, many features scaled linearly with brain volume, with the slope representing the allometric scaling exponent (α; Fig. 2e). Mice exhibited consistently strong allometric relationships with brain volume, characterized by high (absolute) α and *R²* values, indicating that neuronal physiology is tightly coupled to brain growth. In contrast, humans displayed substantially weaker relationships, with lower, typically non-significant α values and low *R²* values for most physiological features (Fig. 2e, Fig. S8, Table S13). These results were robust to extending the human dataset with mixed-origin infant recordings from the first months of life^17^(Fig. S9, Table S12, S14). Thus, where mouse neuronal physiology matures in step with brain growth, human physiology is desynchronized from it. Human neuronal physiology is thus neotenous with respect to the brain’s own protracted growth, maturing on a slower timetable still.

### Protracted functional specialization

Cortical computation, and with it cognition, depends on how individual cortical neurons transform their inputs. In humans, the physiological properties that jointly set this transformation mature over decades (Fig. 1,2), the same timespan over which cognitive development unfolds^3^. We therefore asked whether the neurons present in successive periods of cognitive development are, considered jointly across these properties, distinguishable as populations. To address this, we grouped cells into four age-defined developmental periods corresponding to conventional cognitive stages, infancy (<2 years), childhood (2–12 years), adolescence (12–20 years) and adulthood (>20 years), and compared their multivariate physiology across 21 features spanning subthreshold intrinsic properties, AP kinetics, and firing patterns. Neurons differed significantly between all periods (PERMANOVA: *F*₃,₃₉₈ = 56.7, *R²* = 0.28, *P* = 0.001, n = 403; all pairwise comparisons *P* < 0.01). The physiological profile of human supragranular neurons thus differs from one stage of cognitive development to the next, and so does the transformation they apply to their inputs, measured directly in input-output features withheld from the multivariate analysis (Fig. S11b,c).

To resolve the developmental axis along which neurons mature through this space, and the physiological domains that shape it, we applied principal component analysis to the 21 features. PC1 ordered cells from infancy to adulthood and was loaded most strongly by AP stability and AP waveform, PC2 was dominated by spike-train firing properties, whereas PC3 was loaded most strongly by the subthreshold properties and excitability domains (Fig. 3a,b, Fig. S10a). We then asked whether neurons traverse this space as a single continuum or resolve into distinct states. Unsupervised clustering returned two highly stable clusters (Fig. S10), one spanning the whole of PC1, the other separating along PC2 and appearing only late in development (Fig. 3a-c,d).

**Figure 3.**
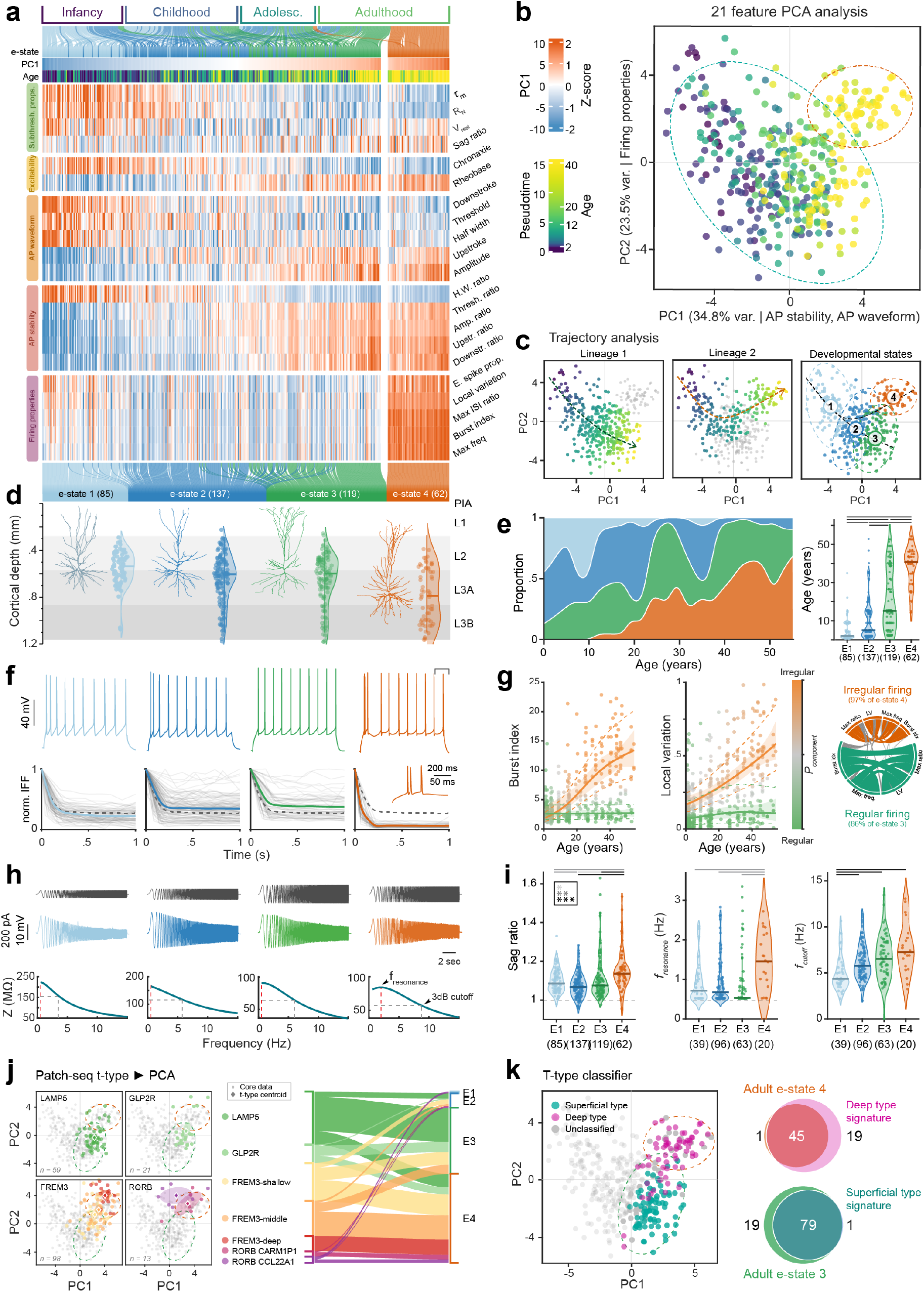
Protracted functional specialization culminates in two adult phenotypes. **a**, Heat map of 21 z-scored features (rows, by domain) for all cells (columns), ordered along PC1 with those of the late-emerging cluster (**b**) at the right; Top alluvials map age categories onto this ordering, coloured by e-state (**c**). **b**, PCA of the same features, cells coloured by age; dashed ellipses show two clusters identified by Gaussian mixture modelling: a developmental continuum (teal) and a late-emerging cluster (orange). **c**, Trajectory analysis, cells coloured by pseudotime along each lineage (grey, not assigned), and the resulting e-states: 1 to 3 partition the continuum by pseudotime, 4 is the late-emerging cluster. **d**, Distribution of e-state neurons across supragranular layers with representative reconstructions. Soma depth differed between e-states (Kruskal-Wallis (KW), *P* = 1.8 × 10⁻⁷), with e-state 1 more superficial and e-state 4 deeper than all other e-states (*P* < 0.01 and *P* < 0.001, resp) **e**, E-state composition of the population with age (left) and age per e-state (right); bars, significant pairwise differences (KW with post hoc tests, *P* < 0.001). **f**, Representative spike trains (top) and normalized instantaneous firing frequency (IFF) of individual cells (grey), with e-state mean (colour) and population mean (dashed). **g**, Mixture models of firing properties against age; points coloured by probability of belonging to either component; lines and shading, component means with 95% c.i.; dashed, ± 1 s.d.. Chord diagram: agreement of component assignment between four firing properties (87–94% concordance). **h**, ZAP stimulus, voltage response and impedance profile per e-state. **i**, Sag ratio, resonance and cutoff frequency per e-state. **j**, AIBS Patch-seq reference data projected into PCA space and alluvial showing assignment of each transcriptomic type to nearest e-state. **k**, Classifier trained on the superficial (LAMP5) and deep (RORB) reference types, applied to the adult e-states 3 and 4. Venn diagrams demonstrate number of cells overlap of e-states 3 and 4 with superficial and deep signatures (16 and 21 unclassified neurons excluded, resp.).

Trajectory analysis resolved two maturational paths through this space. Both share a common origin in infancy and course through childhood before diverging in adolescence to two adult endpoints (Fig. 3c, Fig. S10). We defined four electrophysiological states (e-states) along this path. E-state 1, prevalent in infancy, showed slow AP kinetics and unstable AP waveforms; e-state 2 was the shared intermediate state of childhood and adolescence; and e-states 3 and 4 were the two adult endpoints, both with low membrane resistance, fast time constants and stable APs (Fig. 3a). The e-state composition of the population shifted with age (Fig. 3e, Fig. S10e), and input-output properties including gain, input range and maximum firing rate increased from e-state 1 to both endpoints (Fig. S11d,e).

The two adult states separated along PC2: e-state 3 fired regular spike trains, whereas e-state 4, the late-appearing cluster, initiated firing with bursts of multiple APs (Fig. 3f-g, Fig. S13). Mixture modelling resolved this divergence in firing properties, identifying an irregular-firing component that emerged in adolescence and continued to change into adulthood, to which nearly all e-state 4 neurons belonged (Fig. 3g, Fig. S13d, Table S7). This also reorganises the relationships among features: firing properties and the AP stability and waveform domains together dominate the age-dependent restructuring of feature covariance (Fig. S14). Interestingly, this division into regular-firing and bursting neurons is also present in layer 3 pyramidal neurons of macaque association cortex^13,14^. The supragranular burst firing phenotype thus appears to be a specialization shared with primates, which in humans is established only in adolescence.

In addition to firing properties, e-state 4 was distinguished by a more pronounced voltage sag, elevated resonant and cutoff frequencies and markedly lower impedance across the frequency range tested (Fig. 3h-i, Fig. S12). Together with burst firing, this combination is absent from adult mouse supragranular pyramidal neurons (Fig. 1c, S13d) but characterises the deep supragranular types of adult human^12,24^ and primate^13,15^ cortex, including the RORB-expressing types CARM1P1 and COL22A1, which have no mouse supragranular counterpart and instead resemble infragranular types^12^. These are intratelencephalic projection neurons, and the largest of them, deep FREM3 and CARM1P1, are SMI-32-immunoreactive cells that provide long-range corticocortical projections, as in non-human primates^12^. We therefore tested whether e-state 4 corresponds to these types, using the transcriptomically identified neurons of the human patchseq reference^12^ (Methods). Projected into our PCA space, superficial LAMP5 neurons mapped predominantly to e-state 3 (38 of 59), whereas deep FREM3 and RORB types mapped to e-state 4 (26 of 30; Fig. 3j,k, Fig. S15b). Further, a classifier trained on the superficial and deep poles of the reference taxonomy assigned the deep-type signature to 98% of classified e-state 4 neurons and the superficial-type signature to 81% of classified e-state 3 neurons (Fig. 3k, Fig. S15c).

### A reference atlas of neuronal maturation

Finally, we combined the individual feature trajectories, their variance functions, the age-dependent correlations among them into a single generative model of the developing human supragranular population. Synthetic cohorts can be drawn for any age range, cortical depth and sample size, that reproduce the developmental structure of the recorded population, from the individual trajectories and feature correlations to the divergence into two adult endpoints and the late emergence of e-state 4 (Fig. S16). We make the model available through an online resource, SHERPA (Simulator of Human Electrophysiology, Reference and Population Atlas; https://raimondolab.org/sherpa). SHERPA also serves as a normative reference, in the manner of a growth chart. Datasets too small to estimate the effects of age, depth and variability can draw them from the reference instead, allowing developmental trajectories to be compared and group differences resolved where age and depth may otherwise obscure them (Fig. S17b). Referenced in this way, an independent frontal cortex dataset (192 cells from 44 subjects, 0.1 to 51 years, combining published data^17^ with our own recordings; Table S2) showed no age-dependent departure from the temporal cortex trajectories (Fig. S17a), establishing that the protracted maturation of human supragranular neurons extends beyond temporal cortex to frontal association cortex.

## Discussion

Here we show, with an extensive patch-clamp dataset from developing human and mouse supragranular temporal cortex, that human neurons exhibit a striking neoteny in their physiological development. Where prior rodent and human studies compared physiology between discrete age groups^17,27,28^, we capture the maturation of each physiological feature as a continuous trajectory from infancy to adulthood. Set against the growth of the brain, these trajectories reveal a profound species-specific divergence. In mice, physiological maturation proceeds in step with brain growth, following a strong allometric relationship. In humans the two are desynchronized, with physiological maturation continuing for decades longer, up to seven times more slowly than would be predicted from relative brain growth rates alone, and consistent with extended human cognitive development^3^.

This ontogenetic divergence parallels recent findings that adult human neurons are also outliers in static allometric relationships, exhibiting lower ionic conductances than predicted by their size^29^, so that human neuronal distinctiveness extends across both the developmental and the mature functional domain. Human neoteny has been documented at the cellular and molecular levels, through extended synaptogenesis, delayed myelination and protracted gene expression dynamics^4–8,10^, and its tempo appears to be under cell-intrinsic control^30,31^. Epigenetic barriers constrain the pace of neuronal maturation^9^, and human neurons hold to their schedule even after xenotransplantation into mouse cortex^32^. Our findings establish how far this intrinsic slowing reaches in the intact developing brain, and when the functional properties of cortical neurons mature. Frontal cortex neurons, referenced against the temporal cortex trajectories, follow the same protracted course, showing that our findings generalise beyond temporal cortex to higher association cortex, but whether primary sensory areas mature on the same schedule remains an open question.

Human supragranular neurons mature along a common developmental axis that diverges in adolescence into two adult endpoints, when a subpopulation of neurons (e-state 4) acquires a distinct physiological phenotype that includes burst firing and resonance. These properties are absent from mouse supragranular neurons but characteristic of the SMI-32-positive deep supragranular neurons that send long-range projections to other cortical areas in primates^12^. Bursts are thought to make transmission across long, unreliable projections dependable^33^, to synchronise activity between cortical areas^34^, and to carry information in multiplexed neural codes^35^. The late acquisition of burst firing by the very neurons that carry long-range corticocortical projections, at ages when those projections are still being myelinated^8^, suggests that intrinsic physiology and axonal maturation are jointly timed to bring long-range cortical communication online. Their resonance, in turn, confers frequency selectivity. Since HCN channels also regulate dendritic integration^11,36^ and network rhythmogenesis^37^, their late emergence could endow mature human cortical circuits with integrative capabilities supporting working memory^38,39^.

Long-standing models of postnatal cognitive development have regarded mature information processing as the outcome of synaptic and circuit-level mechanisms^7,40,41^, implicitly assuming that the intrinsic electrical properties of individual neurons remain largely unchanged. In the mouse this is largely true given that intrinsic physiology is established within the first postnatal month, before the protracted period of experience-dependent refinement. In humans, however, the properties that set how a neuron integrates and transforms its inputs are still changing through adolescence and into adulthood. These are the properties to which distinctive computational capabilities of human pyramidal neurons have been attributed, from subthreshold features that shape integration, charge transfer^42,43^ and theta-band resonance^11^, to maintaining the fast action potential kinetics that support high-bandwidth encoding^19,44^ and correlate with cognitive ability^20,21^. Whilst synaptic weights and connectivity determine which inputs a neuron receives, the transformation it applies to them is set by the properties reported here. As studies in well-characterised circuits show^45,46^, the capacity of synaptic mechanisms to compensate for changes in intrinsic properties is bounded. Therefore, the maturation of human cortical circuits is best understood as an interplay between intrinsic properties and synaptic plasticity. Weighing the relative importance of these two contributions will require modelling with realistic neuronal populations in development, which the joint developmental model presented here and released through SHERPA now supplies.

The systematic decrease in physiological variability with age in both species, driven in humans by reductions in within-subject variance, suggests a progression from plastic, variable neuronal states toward stable, specialized profiles. The juvenile properties we identified in humans, high input resistance, long membrane time constants and slower action potentials, make neurons more excitable and responsive to small inputs, but with limited speed or temporal resolution. Analogous to dropout in artificial neural networks^47^, or to the annealing schedule by which optimization algorithms escape local minima before settling^48^, this early variability might prevent overfitting during critical developmental learning periods and promote robust generalization later in life. As development progresses, membranes become leakier and faster, action potentials narrow and accelerate, and rheobase increases, converting neurons from excitable but imprecise elements into selective, rapidly responding units capable of sustained high-frequency firing with reliable timing^19,44^. This trajectory likely optimizes cortical circuits first for maximal plasticity and broad learning during childhood, and thereafter for the stability and precision required for reliable information processing, mature executive function and higher-order cognition^49^.

The extended timeline of human neuronal maturation coincides with the progressive acquisition of complex cognitive functions, and perturbations of this schedule may seed neurodevelopmental disorders^50^. Premature maturation could truncate periods of heightened plasticity, delayed maturation might impair the precise temporal dynamics needed for higher cognition, and disorders that surface only later in life may stem from latent timing irregularities that escape detection until circuits demand fully mature properties. Studying such disorders in rodents therefore requires care. Much rodent electrophysiology is performed on two-week-old animals^51^, an age corresponding to middle childhood in humans for many physiological features. Our feature-by-feature age mapping provides a means of selecting age-appropriate models for specific human developmental periods, and offers a starting point for relating the mouse developmental literature to a human timescale from the perspective of cortical neuronal physiology, rather than through a single translation anchored on milestones such as brain growth, weaning or sexual maturity. In humans, SHERPA provides the normative reference, in the manner of a growth chart, that gives even the small datasets clinical tissue yields the sensitivity to reveal a departure from the expected developmental course. Developmental timing thus joins circuit topology and cellular heterogeneity as a fundamental principle shaping the human brain, one that may help account for when cognitive capacities emerge and, when perturbed, for the characteristic ages at which developmental brain disorders arise.

## Acknowledgements

We would like to thank patients, their parents and families for the generous donation of resected tissue for research purposes. We thank hospital clinical and support staff including Kelsey Bester, Lucretia van der Horst, Jodi Jacobs and Natalie Vincent for logistical support as well as members of the Raimondo Lab, Dorit Hockman, Rachael Dangarembizi, Christopher Dulla and Colin Akerman for advice and comments. We are grateful to Simon Weiler and Anne-Marie Oswald for sharing data from previously published mouse datasets. We thank Henry Markram (École Polytechnique Fédérale de Lausanne) and Tobias Boenhoffer (Max Planck Institute for Biological Intelligence) for generous donation of patch-clamp equipment to the UCT, Rodrigo Perrin, Charles Harris and Volker Staiger for assistance setting up of Ephys recording equipment and software, Caron Jacobs and the Africa Microscopy Initiative Imaging Centre (RRID: SCR_025881) for microscopy support and the UCT ICTS High Performance Computing team for support using the high performance cluster. This research received support from a European Molecular Biology Organisation Fellowship (EMBO ALTF 415-2018; MBV), a UCT Faculty of Health Sciences Fellowship (MBV), a Claude Leon Foundation Fellowship (MBV), a Royal Society Newton Advanced Fellowship (JVR), the SA National Research Foundation (JVR), the Blue Brain Project (JVR), the Gabriel Foundation (JVR), a Wellcome Trust Seed Award (214042/Z/18/Z) (JVR), the South African Medical Research Council (JVR) and the FLAIR Fellowship Programme (FLR\R1\190829) (JVR): a partnership between the African Academy of Sciences and the Royal Society funded by the UK Government’s Global Challenges Research Fund and a Wellcome Trust International Intermediate Fellowship (222968/Z/21/Z) (JVR).

## Methods

### Human and mouse brain samples

Human temporal cortex tissue was obtained from 80 donors (37 females, 43 males; 0.9-54.4 years) who underwent surgical treatment for epilepsy with various etiologies (Table S1). Tissue originated from the inferior, medial and superior gyrus of the temporal lobe, approximately 2–6 cm posterior to the temporal pole and was removed for access to deeper brain structures, or comprised the least affected, most normal tissue surrounding the primary affected area, lesion or tumor. The cohort comprised 34 donors from South Africa of diverse population groups: 18 black african, 7 mixed ancestry, and 9 white/caucasian individuals. The other 46 donors were from the Netherlands, for which population information was unavailable. Genetic ancestry was not assessed. These descriptors serve to illustrate the inclusion of underrepresented populations in our study sample, and play no role in participant recruitment, study design, analysis, or interpretation of results. All procedures on human tissue were performed with the approval of the Human Research Ethical Committee of the University of Cape Town and the Medical Ethical Committee of the VU University Medical Centre. Written informed consent was obtained prior to surgery from all patients, or in case of minors, from their guardians.

Mouse brain samples were obtained from C57BL/6J mice, covering the full postnatal developmental period from the first postnatal day to adulthood (P0.86-P86, 42 animals), following anaesthetized decapitation. All procedures for mouse neocortical brain slice preparation were approved by the Animal Research Ethics Committee of the University of Cape Town.

### Brain slice preparation

Following resection (humans) or extraction of the brain (mouse), tissue was transferred to ice-cold artificial cerebrospinal fluid (aCSF) containing (in mM): 110 choline chloride, 26 NaHCO₃, 10 D-glucose, 11.6 sodium ascorbate, 7 MgCl₂, 3.1 sodium pyruvate, 2.5 KCl, 1.25 NaH₂PO₄, and 0.5 CaCl₂, and transported to the laboratory within 10-20 minutes. Tissue blocks were embedded in low melting point agar for sectioning (350 μm) on a Compresstome (Precisionary Instruments) or processed for vibratome slicing as previously described^52,53^. Slices were stored at 34°C for 15-30 minutes, then allowed to recover for at least 1 hour at room temperature before recording in aCSF containing (in mM): 125 NaCl, 3 KCl, 1.25 NaH₂PO₄, 1 MgSO₄, 2 CaCl₂, 26 NaHCO₃, and 10 D-glucose. All solutions were continuously bubbled with carbogen (95% O₂, 5% CO₂) and maintained at 300 mOsm osmolarity. Mouse brain slices were prepared using identical procedures, with coronal sections including temporal association cortex.

### Characterization of neuronal physiology

Following recovery, slices were placed in a recording chamber and perfused with aCSF (3–4 ml/min, 31–34 °C). Pyramidal neurons in supragranular layers 2-3 were identified with oblique illumination or differential interference contrast microscopy. Whole-cell patch-clamp recordings were then made using borosilicate glass pipettes with fire-polished tips (4.0–6.0MΩ resistance) filled with intracellular solution containing (mM): 110 K-gluconate; 10 KCl; 10 HEPES; 10 K2Phosphocreatine; 4 ATP-Mg; 0.4 GTP, biocytin 5mg/ml (pH adjusted with KOH to 7.3; 280–290 mOsm). Whole-cell patch-clamp recordings were made using Axopatch 200B or MultiClamp 700 A/B amplifiers (Axon Instruments), sampling at 10-50 kHz and low pass filtering at 5-10 kHz. Recordings were digitized with InstruTech ITC-1600 or Axon Digidata 1440A digitizers and acquired using PulseQ (v1.01, Funetics), WinWCP (v5.2.4, University of Strathclyde) or pClamp (Axon Instruments) acquisition software.

### Morphology & Histology

After recordings, slices were fixed in 4% paraformaldehyde overnight, rinsed with phosphate buffer (PB) and stored in PB-azide at 4°C. Biocytin-filled cells were visualized using either the avidin– biotin–peroxidase method^54^ and mounted on slides and embedded in Mowiol, or by Alexa488-streptavidin (1:500, Thermo Fisher Scientific) for 48 hours at room temperature, followed by 3 washes in PB before being mounted in Vectashield (Vector Laboratories).

For histological assessment, recorded slices were co-stained with DAPI (4’,6-diamidino-2-phenylindole, 5 µM) for 20 minutes at room temperature to visualize cell nuclei and cortical architecture and imaged on a Zeiss Axioscan 7 automated fluorescence slide scanner at 20x. Layer and cortical thickness was subsequently measured using ImageJ^55^. The distance of retrieved somata to pia was measured along the apical dendrite, and position within L2-3 was verified (Fig S1). For neurons not retrieved in post-hoc staining, the pia-soma distance measured during recording was used. Only neurons <1500 µm from pia were included in this study (range: 237-1500 µm, mean 765 µm).

A subset of cells was selected for full 3D reconstruction of neuronal morphology using Neurolucida software (MBF Bioscience). Z-stack mosaics were acquired at 0.5 μm intervals on a Zeiss LSM 880 Airyscan confocal microscope (Carl Zeiss, hosted by UCT Human Biology Dept.) or an LSM 980 Airyscan 2 (hosted by the African Microscopy Initiative (AMI)), using a 63× 1.4 NA oil-immersion objective (Fig. 3d, Fig. S1).

### Electrophysiological feature extraction & analysis

#### Stimulation protocols

Recorded neurons were subjected to a suite of current clamp injection protocols to characterize a wide range of physiological properties. These included current injection steps, ramps and chirps of varying length and amplitude. The intensity of current stimuli was adjusted for each neuron, uniformly scaling protocols up or down to ensure neurons were driven to a similar degree regardless of input resistance. Neurons with resting membrane potential (*Vres*t) close to firing threshold were hyperpolarized to -60 mV with a holding current to facilitate physiological characterization. Recordings with access resistance *Ra* > 30MΩ, or with unstable resting membrane potential were discarded. Data files were then analyzed using custom MATLAB scripts. All sweeps were visually inspected and cases where events (e.g. noise, spontaneous activity, sudden change in *Ra*) challenged the normal extraction of physiological features were excluded.

#### Subthreshold properties

Subthreshold membrane properties were calculated as follows^11^ (Fig. S2): Input resistance *RN* was the *I-V* slope of a linear fit to the minimum voltage deflection measured within 80 ms of onset of hyperpolarizing current injection (0.5-1s). Membrane time constant was the average time constant of single exponential fits (*R^2^* > 0.9) to the same hyperpolarizing current pulses. To obtain sag ratio, *RN* was divided by resistance at steady state (*RSS*, where both fits had *R^2^* > 0.8). Resting membrane potential (*Vrest*) was measured shortly after whole-cell configuration. In cases where hyperpolarising holding current was applied to prevent firing, Vrest was calculated from injected holding current and input resistance: *V_rest_*= *V_recorded_*+ (*I*_ℎ*old*_ × *R*_N_).

#### Excitability properties

Rheobase (*Rb*) and chronaxie (*Cx*) were estimated fitting the linear charge-delay relationship: *Q* = *Rb* × (*delay* + *Cx*)^56,57^, where charge *Q* was the product of AP delay and injected current (Fig. S3). This approach allowed estimation or rheobase (slope) and chronaxie (-x intercept) from a diverse set of protocols or from cells where a limited number of suprathreshold responses were available (required: min 3 ms stimulus duration, ≥3 current levels, ≥3 sweeps, max 4 mV baseline change between sweeps and fit *R^2^* > 0.75). In a small subset of cells where fitting was not possible, rheobase was based on the minimum step current injection (≥500 ms) to elicit an AP.

#### Action potential waveform

Action potentials (APs) were detected as peaks exceeding 0 mV and threshold was defined as the voltage at which the derivative was last <15 mV/ms before the peak (Fig. S4). Amplitude was the difference between threshold and peak voltage, and AP half-width measured as the ms time between AP rise and fall at *Vthresh* + ½ AP amplitude. Cells with half-width <0.5 ms were excluded as presumed interneurons. AP upstroke and downstroke were the max and minimum values of the AP derivative (mV/ms) during the rising and falling phase of the AP, the upstroke-downstroke ratio was the absolute ratio of the two. The instantaneous firing frequency in Hz was *f_inst_* = 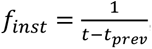, where *tprev* was the time of last spike.

The total pool of recorded APs exceeded 100,000. Although our AP detection and feature calculation algorithms performed well across ages and firing intensities, at high firing rates where neurons can no longer sustain their AP train, the reliability of AP detection methods can be compromised. We hence imposed broad quality control constraints for AP threshold (between -60 and 0 mV), amplitude (20-120 mV), half-width (0.5-10 ms), upstroke (0-800 mV/ms), downstroke (-250-0 mV/ms), and *finst* (<200 Hz). APs failing any of these criteria were excluded from analyses of AP kinetics, in addition to those recorded during high-intensity current injections aimed at pushing neurons to their maximum firing rate. A few cells were observed to receive presumably disynaptic inhibitory synaptic input following an action potential. In such cases, kinetics of APs later in the train were compromised and therefore excluded from analysis.

#### Action potential stability

Susceptibility to use-dependent change of the 5 AP waveform metrics was assessed by dividing the values of 5^th^-10^th^ AP by the values of the 1^st^ APs in a train, using APs with *finst* <20 Hz (Fig. S4). The AP stability index was the first PCA component on these five standardised AP ratio metrics, which accounted for 88% and 72% of variance in human and mouse, respectively, which was linearly rescaled so that no change across the train scored 1 and complete collapse scored 0 (Fig. S4e).

#### Firing properties

Firing patterns were analyzed using the inter-spike intervals (ISI) of trains of APs (7-13 spikes) evoked by 1s depolarizing current injections. Measures selected to capture distinct aspects of spike train regularity and temporal structure were: (1) Burst index, defined as the ratio of the last ISI to the first ISI, quantifies the degree of spike frequency adaptation within a train. (2) Local variation (LV), defined as: 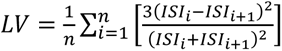, where the summation is over all pairs (i,j) of ISI in the spike train, and n is the total number of ISI pairs. This approach quantifies the irregularity of spike timing by measuring how much consecutive intervals vary relative to their sum, with higher values indicating greater variability. (3) Maximum instantaneous frequency (*fmax.inst*), calculated as the inverse of the smallest ISI (Hz), captures peak firing frequency within the train. (4) Maximum ISI ratio, defined as the ratio of the largest to smallest ISI, quantifies the range of temporal variability. (5) Early spike proportion was the fraction of action potentials occurring within the first 100 ms of stimulus onset, reflecting the temporal distribution of spiking activity.

#### Input-output features

Gain was defined as the slope of the linear portion of the *f-I* (frequency-current) relationship. Full *f-I* curves were fit using the Hill function: *F*(*I*) = 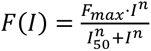, where *F_max_* is maximum firing rate, *F_50_* the current for half-maximum response, and *n* the Hill coefficient controlling curve steepness. Only neurons that were pushed to fire close to predicted maximum were included. Input range was defined as the current span between levels that produce 10-90% of *Fmax*.

Subthreshold filter properties were assessed using a chirp current injection protocol that consisted of a constant amplitude sinusoidal current stimulus with linearly increasing frequency over time (0.5-15 Hz over 15-30 s). The impedance amplitude profile (ZAP) was computed by taking the ratio of the fast Fourier transform of the recorded voltage response to the fast Fourier transform of the injected current waveform. Resonant frequency was identified as the frequency corresponding to maximum impedance (*Zmax*) in the ZAP profile. The 3 dB bandwidth cutoff frequency was calculated as: 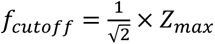

### Datasets

In total, the final human temporal cortex dataset consisted of 584 recordings of supragranular layer 2/3 pyramidal neurons from the temporal cortex of 80 donors aged 0.91 to 54.4 years (37 female, 43 male; Table S1-2). The parallel mouse dataset consisted of 151 layer 2/3 pyramidal neurons from the temporal association cortex of 42 C57BL/6J wild-type animals aged P0.86 to P86. For a set of supplementary analyses we assembled further human and mouse developmental datasets of layer 2/3 pyramidal neuron physiology, combining our own recordings with published data. These include a human frontal cortex dataset^17,58^, an extended infant dataset including recordings from even younger cases of mixed cortical origin^17^, and mouse frontal, auditory and visual cortex datasets^28,59,60^ (Table S2). For all but mouse auditory cortex, raw recordings were obtained from the authors or public repositories and processed through the same MATLAB feature extraction and quality control procedures as our own recordings.

Human recordings were digitized at a range of sampling intervals, which may impact the precision with which fast events like APs are captured. To detect sampling frequency-related bias in action potential kinetics, we compared measurements from the same human neurons (n=150) recorded at high and low sampling frequencies using Bland-Altman analysis. Most AP features showed <5% difference in values between sampling regimes and measurements were therefore pooled, but AP upstroke and downstroke velocities showed systematic bias >10%. This was corrected by fitting a first and second-degree polynomial, respectively, to the high-vs-low sampled values, which was then used to correct the values derived from low frequency acquisition.

### Generalised additive mixed models

Developmental trajectories of physiological features were modelled as generalized additive mixed models (GAMMs^61^), fitted in R (v4.5.2) using the glmmTMB package (v1.1.14)^62,63^. A total of 44 models (22 per species, Tables S3-10) were designed following a stepwise approach, starting with the simplest specification and progressively adding complexity, using likelihood-ratio tests to guide selection of the most parsimonious model that best captured the developmental trajectory. Age-dependent change was first modelled as a linear effect and was then entered as a natural cubic spline with 2 or 3 degrees of freedom, allowing trajectories to take a non-linear form. Soma depth, and sex in the human models, were then evaluated as covariates, both as a main effect and in interaction with the age spline. All GAMMs included a subject-level random intercept to account for sampling multiple cells per subject. For AP waveform features, AP number (1,10) was included as an additional fixed effect in interaction with the age spline, with cells nested within the subject random effect. Human firing properties were fitted as a two-component mixture models (GAMLSS, gamlssMX^64,65^), in which each component carries its own mean trajectory and variance and the mixing proportion is itself modelled as a function of age, and was compared with the single-component model of the same specification by BIC. A two-component model was accepted if ΔBIC>25 (Fig. S13), after which each cell was assigned to the regular or the irregular component by its posterior probability.

The t-family distribution was chosen for its robustness to minor deviations from normality that are common in electrophysiological data. Heteroskedasticity was addressed by incorporating dispersion formulas to allow residual variance to change with age, testing both linear and spline-based dispersion models. Final models (Table S3-10) were evaluated using the DHARMa package model diagnostics^66^. For a subset of models, diagnostics showed minor deviations in the upper or lower residual quantiles, which was deemed acceptable given our focus on mean developmental trends and variance changes. For human AP downstroke, a simpler model with better residual diagnostics was adopted. For chronaxie and mouse AP stability, AIC favoured a more complex model and although the improvement was not significant (*P* = .06 to .10), we adopted them for consistency with other features of the species.

Predictive accuracy was assessed by leave-one-out cross-validation, in which each cell’s value was predicted from a model refitted with that cell held out, given only the age of its parent subject and its soma depth. The median cross-validated R*²* was 0.19 in human and 0.74 in mouse, and the residuals showed no systematic trend with age (Fig. S6b,c, Table S11). The models presented here therefore account for the hierarchical structure of our data, including non-independence of observations from the same subject, depth-dependent effects within the supragranular layers, and age-related changes in variance, all critical factors for accurate characterization of neuronal maturation trajectories.

GAMM mean predictions and confidence intervals were derived from a parametric bootstrap procedure, where 10,000 simulated datasets were generated from the fitted model parameters and re-fitted employing the University of Cape Town’s ICTS High Performance Computing Cluster. From each fit, estimated marginal means (EMMs) were extracted using the sjPlot package. Layer-specific predictions were derived as partial effects by fixing cortical depth to representative values for each layer (L2: 400μm, L3A: 700μm, L3B: 1000μm, L3C: 1300μm). Similarly, female- or male-specific partial effects were obtained from models where biological sex was included. Variance predictions were computed by applying the model’s dispersion formula (Table S3). Predictions were spline-interpolated across the age range, with mean and 95% confidence intervals calculated from the resulting distribution. Parametric bootstrapping provides more robust and accurate confidence intervals compared to direct model predictions, particularly for mixed-effects models like these with non-Gaussian distributions and heteroskedastic variance structures.

### Sensitivity analyses

#### Leave-one-subject-out analysis

To assess the stability of the fitted trajectories, and to establish that they are not shaped by individual subjects, each human model was refitted with one subject omitted in turn, and the refitted trajectories were compared with the fit to the full dataset (Fig. S6a). The trajectories were essentially unchanged throughout.

#### Sampled age range and model flexibility

Our temporal cortex recordings start at 10 months. A recent report describes considerable change in human supragranular neurons over the first year of life^17^. To resolve how change in this early period, together with the flexibility of our model specification may bear on the estimated trajectories, we assembled an extended infant dataset of mixed cortical origin: raw recordings from Barzó et al. were combined with infant recordings of our own and processed through the same feature extraction and quality control procedures as the core dataset (73 cells | 9 donors; 0.08-1.29 years; Table S2). Missing soma depths (n = 36) were imputed by extended data mean. Select features were then refitted using this extended dataset, using models of increasing flexibility with 2 and 3 degrees of freedom, compared by likelihood-ratio test and BIC as above (Table S12). To establish that the shape of the trajectories was not constrained by the spline specification, the features were also fitted with a penalised spline (GAMLSS) carrying the same fixed and random effects, dispersion model and response distribution, and differing only in how the smooth is built: knots were placed densely across the age range, including one at 1 year, and the roughness penalty was set so that the fitted smooth retained five effective degrees of freedom, intentionally more flexibility than the data support. Each refitted model was carried through the downstream analyses, and the resulting trajectories, maturation peak ages, dynamic time warping scale factors and allometric scaling exponents were compared with those of the main models (Fig. S9, Tables S12 and S14).

#### Epilepsy etiology

To ensure that our developmental findings reflected normal cortical maturation rather than epilepsy-related pathology, we assessed whether epilepsy etiology or seizure history influenced the neuronal properties and developmental trajectories we observed. To assess the potential effects of epilepsy etiology on cortical lamination, we fit GAMMs to analyze cortical thickness and supragranular layer 2-3 thickness as functions of age and etiology groups (*t*ℎ*ickness* ∼ *age* + *etiology*_*group*; Table S1). Cortical thickness showed a significant age-related decrease consistent with previous MRI-based reports, but no significant differences between epilepsy etiologies (Fig. S1d). Supragranular layer 2-3 thickness remained stable across age and also showed no significant differences between epilepsy etiologies (Fig. S1e). These findings indicate that the diverse epilepsy etiologies in our sample did not affect cortical architecture, which is in line with the apparently normal lamination and appearance of neurons during live inspection of brain slices prior to recording.

The influence of epilepsy etiology and seizure history on neuronal physiological properties and their developmental trajectories was assessed by including these factors in the GAMMs for each physiological feature. Patient etiologies were grouped into four broad categories: Developmental (FCD, HME, NDD), Hippocampal sclerosis (MTS), Structural (SVM, LGT), and Other/Unknown (non-lesional, unavailable etiology). To address collinearity between epilepsy duration and age (older patients will tend to have longer epilepsy exposure), we calculated age-adjusted epilepsy residuals using the formula: residuals = YearsEpilepsy - f(age), where f(age) represents a natural spline fit of epilepsy duration on age. These residuals quantify seizure exposure independent of subject age. We then tested three models: (1) addition of etiology group as a categorical fixed effect, (2) addition of age-adjusted epilepsy residuals as a continuous predictor, and (3) inclusion of both epilepsy residuals and their interaction with age to test whether seizure sensitivity varies across development, each compared with the base model by likelihood-ratio test (Table S15). Epilepsy duration had no detectable effect on any feature, either as a main effect or in interaction with age. Etiology group improved the fit for two of the twelve features, sag ratio and AP stability. The effects were small and BIC did not support including the term in either model (ΔBIC +5.7 and +7.5, Table S15). Overall, epilepsy-related factors did not substantially influence the developmental trajectories of neuronal physiological properties, in line with earlier reports that the impact of patient disease history on neuronal physiology is minimal^12,53^, but see^23^. Together, this analysis adds to the growing evidence that surgically resected tissue as used in the present study shows no measurable effects of patient disease history on cortical lamination^21^, pathological marker expression^12,67^, neuronal physiology^12,21,53^, morphology^68^, or gene transcription^12,69^.

### Variance components

Some features were log-transformed prior to fitting (Table S3). For display/interpretability purposes and for normalization of developmental trajectories (below), mean predictions of log-transformed variables were back-transformed to the original scale by exponentiation. Variance estimates were approximated on the original scale using the log-normal transformation formula *exp*(2 ∗ *μ*) × (*exp*(*σ*^2^) − 1), where *μ* and *σ²* represent the predicted mean and variance on the log scale, respectively.

To estimate within-subject (*σ²within*) and between-subject (*σ²between*) variance components across age categories, we again used parametric bootstrap simulations from the final GAMMs for each feature. Here and in following analyses, simulations were generated with soma depth fixed at 750 μm (approx. L3A and close to mean depth of our sample), to eliminate variability due to differential depth sampling. For each replicate, variance components and intraclass correlation coefficients 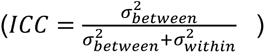 were calculated per age category (Fig. S2g,h, S3f,g). Variance components were estimated by one-way random-effects analysis of variance, with subject as the grouping factor. We estimated *σ²within* as the within-subject mean square. The variance of subject means overestimates the between-subject variance, because each subject mean also carries sampling error. We therefore estimated σ²*between* as (*MSbetween* − *σ²within*)/n₀. Here, *MSbetween* is the between-subject mean square and 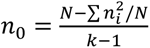 is the effective number of cells per subject, where *N* is the number of cells, *k* the number of subjects, and *ni* the number of cells from subject *i* within the age category. Negative estimates were set to zero. Pairwise differences in variance components and ICCs between age categories were considered significant if the Bonferroni-adjusted confidence intervals from the bootstrap distributions excluded zero.

### Maturation peaks

The age of reaching adult maturity was estimated using the bootstrap replicates for each physiological feature that exhibited significant age-dependent change (Fig. 2a). For each replicate, adult ‘peak’ age was defined as the first time beyond the juvenile period (>12 years human, >12 days mouse) where developmental change slowed to <5% of the maximum rate of change (Fig. S7a). For some non-monotonic trajectories, the identified plateau value was inconsistent with the overall developmental direction (e.g. in case of saddle points, where trajectories increased overall but showed local minima, or vice versa). In such cases, only plateau values >20 were retained. Adult maturity age was defined as the median of the resulting adult peak age distribution. The associated uncertainty was quantified by the IQR and 2.5-97.5% percentiles, which reflect how parameter estimation uncertainty in the fitted model translates into uncertainty about maturation timing. To align human and mouse age axes (Fig. 1d), respective feature trajectories were expressed relative to their own adult values and aligned using least squares to identify the same relative range of change.

The average developmental trajectory of each feature was normalized to a 0-1 scale based on the parameter’s value at the youngest age sampled and the value at its identified adult age: *N*(*t*) = 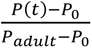 for increasing trajectories and 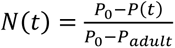 for decreasing trajectories, where *P(t)* is the parameter value at age *t*, *P₀* is the initial/infant value, and *Padult* is the adult value. For cross-species comparison via dynamic time warping analysis (below), mouse trajectories were normalized over an equivalent neurodevelopmental window: mouse *P₀* was redefined as the parameter value at postnatal day P6.3, the age when mouse brain volume reaches 57% of adult volume, matching the relative brain maturation stage of our youngest human subject (0.91 years). The same threshold was applied when displaying mouse developmental trajectories: in Fig. 1c mouse data are shown from P6.3 onward. Mouse ages below this threshold are conventionally taken to correspond to prenatal stages of human development^70^.

### Brain development curves

We obtained postnatal brain development curves for both species from published literature that characterized the postnatal growth of human^71^ and mouse^72^ brain structures using MRI. The human brain volume developmental trajectory was taken from the lifespan brain charts of Bethlehem et al. (2022), where the fitted population-median trajectory for total cerebral volume was interpolated onto a 0.1 year grid using cubic interpolation. Mouse brain volume trajectories were modeled using logistic growth curves^72^: 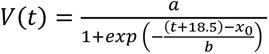, where *t* is postnatal age in days. Mouse cerebral volume was calculated by subtracting cerebellar from total brain volume, obtained using the parameters for total brain (*a* = 429, *x*_0_ = 24.0, *b* = 4.2) and cerebellum volume growth (*a* = 48, *x*_0_ = 27.0, *b* = 3.6)^72^. For the dynamic time warping analysis procedures below, brain growth curves were normalized as described for physiological maturation trajectories above.

### Dynamic time warping analysis

For cross-species comparison of brain anatomical and neuron physiological maturation rates, we performed dynamic time warping (DTW) analysis on normalized developmental trajectories for human and mouse. The DTW algorithm constructs a pairwise distance matrix between all time points of the two developmental series (days for mouse, years for humans), then identifies the optimal warping path that minimizes the cumulative distance between aligned ages (Fig. S7b,c). Both species exhibited developmental trajectories with similar shapes typical of biological maturation processes, resulting in nearly linear optimal warping paths. This linear relationship indicates that relative maturation rates between species can be effectively captured by a single scaling factor, the slope of the warping path, which quantifies the temporal scaling of human and mouse development. Only features that had a significant age effect in the GAMMs for both species were included in this analysis. For brain development, the trajectories spanned from birth to when 99% brain volume was reached in both species. For physiological features, trajectories were normalized over an equivalent brain growth window (57-99%, see above). Developmental scaling coefficients were derived from the ratio of physiological to anatomical scale factors (physiology/brain growth), capturing the extent to which physiological and anatomical maturation are proportionally scaled across species.

### Allometry

To explore allometric relationships between maturation of physiological features and brain volume growth, we used all data acquired up to peak brain volume (12.6 years in human, P34 in mouse), or up to the adult peak age of the feature, whichever came first. Feature values and brain volumes were log-transformed and fitted with a linear mixed model, log10(feature) ∼ log10(brain volume) + soma depth + (1 | subject), in which the slope on brain volume represents the allometric scaling exponent α in the power law function *y* = *bx^α^*. Statistical significance of allometric scaling differences between species was assessed with a two-sided Wald test on the difference between the human and mouse estimates of α (Table S13).

### Multivariate analysis and developmental staging

Multivariate analyses were performed on a dataset of 403 neurons (≤1200 μm) with no missing values for 21 electrophysiological features, which included all sub- and suprathreshold properties, AP kinetics and stability measures mentioned above, with the addition of a set of spike-train regularity measures. Variables were transformed where necessary (Table S3), standardized, and PCA analysis was performed. The first 3 components carried most of the developmental signal and were used for PERMANOVA and further clustering. PERMANOVA (vegan adonis2, v2.7-3) tested the effect of developmental period on Euclidean distances between cells. Soma depth was entered as a covariate before developmental period, and significance was assessed with 999 permutations. Multivariate dispersion did not differ between periods (PERMDISP with bias adjustment, 999 permutations; *P* = 0.09). Pairwise tests were done with Bonferroni correction for the six comparisons.

Gaussian mixture models (mclust version 6.1.2^73^, VVI covariance) were then fitted over 1-5 components, choosing the optimum of 3 components at the elbow of the BIC curve. These were then merged using the entropy criterion of clustCombi^74^, which combines components that subdivide a continuum rather than separate distinct groups. The merged solution comprised two superclusters: a developmental continuum ordered along the first principal component, and a separate cluster emerging late in development (Fig. 3a-e, Fig. S10). Stability was assessed by resampling cells with replacement and re-running the entire chain on each of 1,000 draws, comprising rescaling, principal component analysis, mixture fitting, component selection and entropy merging. Each draw’s partition was compared with the reference partition by the Jaccard index of the matched sets. Two superclusters were recovered in 96.5% of draws, with a median Jaccard index of 0.995 for the developmental continuum and 0.975 for the late-emerging cluster (Fig. S10c).

To order cells along the developmental axis through the space of physiological properties, we employed trajectory inference analysis using the Slingshot package^75^. Principal curves were fitted with in PC space from the youngest to the two oldest components, yielding two lineages that shared a common course from infancy and diverging towards the two adult endpoints. Pseudotime represents each cell’s relative position along the inferred developmental trajectory and as used to define four electrophysiological states (e-states). The late-emerging cluster forms e-state 4. The developmental continuum was divided into three sequential e-states by pseudotime rank (0.25 and 0.65 quantiles). State membership was assessed under the same resampling procedure, with trajectory fitting and staging repeated in full on each draw, and reproduced with median Jaccard indices of 0.895, 0.825 and 0.871 for e-states 1 to 3 and 0.975 for e-state 4 (Fig. S10c). Importantly, we do not treat these as discrete developmental classes; they serve as reference points along the maturational axis, against which features held out from the clustering can be compared.

### Patch-seq mapping and classification

Patch-seq recordings from Berg et al. (2021) were processed through our own pipeline and the 21 clustering features extracted. For the projection, each feature was aligned to our adult recordings by median and interquartile range on the scale at which it was modelled (Fig. S15a), with the alignment estimated on the reference cells within our depth range (≤1200 μm); missing chronaxie and rheobase values were imputed by within-type median. Aligned cells were carried through the same scaling and principal component rotation as our own, and e-state assigned by nearest centroid (Fig. 3j, S15b).

A random forest (randomForestSRC, 1000 trees) was trained on LAMP5 and RORB types, the two poles of the reference taxonomy. The classifier used 19 of the 21 features, with chronaxie and local variation excluded (chronaxie fits were poorly constrained or missing in Patch-seq data, and local variation correlated differently with other features in the two datasets). Each dataset was transformed and z-scored to itself, and the classifier fitted in the first three principal components of a pooled decomposition of reference and query cells, a truncation in which dataset of origin is no longer recoverable. Out-of-bag accuracy was 0.92, and a cell was classified only where at least two thirds of trees agreed. The two training types differ in laminar position as well as transcriptomic identity. To verify the classifier encoded more than soma depth, we compared linear models of the score on depth with and without transcriptomic type. Within the reference neurons, over the depth range where FREM3 and RORB overlap (749 to 1,455 μm, 41 cells each), transcriptomic type accounted for variation in the score that depth could not explain (*F*(1, 79) = 14.2, *P* = 3.1 × 10⁻⁴). Applied to our adult cells, the score separated e-states 3 and 4 better than measured depth did (AUC 0.91 against 0.67), and in a joint logistic model it remained strongly predictive after adjustment for depth (odds ratio 9.3 per s.d., *P* = 1.5 × 10⁻¹¹) while depth added nothing detectable once the score was included (*P* = 0.14). We therefore refer to its output as a deep-type signature score (Fig. 3k, S15c).

### Joint developmental model and population simulation

For each species, the 21 models describing maturation trajectories across physiological domains were combined into a single generative model of the supragranular population (Fig. S16). The model draws new synthetic populations of any specified hierarchy and distribution: the number of subjects, the cells per subject, the subject ages and the soma depths. The draw preserves the hierarchical structure of the data, with cells nested in subjects. For each feature, simulated values follow its fitted trajectory at the cell’s age and soma depth, with a spread that combines the model’s between-subject and within-subject variance. Dependence between features is imposed by a Gaussian copula, estimated on the recorded cells after converting each value to a standard normal score with its fitted model. For a given feature, the score of cell *i* from subject *j* is zij = √ICC · uj + √(1 − ICC) · eij, where *uj* is a subject term, drawn once per subject and shared by all cells of that subject, and *eij* is a cell term, drawn once per cell. Each feature has one ICC for all ages, estimated by restricted maximum likelihood (REML) from a random-intercept model fitted to its scores. Correlations between features were estimated at two levels. Between subjects, they were estimated from subject averages, pooled over all ages. Within subjects, they were estimated from each cell’s deviation from its subject’s average, separately for each age category. To stabilise matrices with a smaller number of cells, the weight on a category’s own matrix was shrunk toward the matrix pooled over all ages (n/(n + 60), where n is its number of cells). For human neurons, the procedure has two stages. First, the 16 subthreshold, excitability, AP waveform and AP stability features are drawn in one copula step that includes the subject terms. Second, each cell is assigned to the regular or the irregular firing component by a Bernoulli draw, *P(irregular) ∼ ns(age, 2) + soma depth + RN*, and its five firing properties are drawn in a second copula step coupled to the first-stage features. Values were capped at the observed range of each feature extended by 50% at either end and prevented from crossing zero where the reference data are strictly signed, which affects fewer than 0.2% of cells. For mouse, all features are drawn in one step, with correlations pooled across ages. Fidelity was assessed on 1,000 cohorts matched to the recorded dataset subject by subject, in age, number of cells and sampled depths, each passed through the clustering and staging procedure above (Fig. S16c-e).

### Normative referencing

External smaller datasets can be referenced against the GAMMs presented here in the manner of a growth chart. Each cell is scored as a centile, with the effects of age and soma depth and the expected variance taken from the reference, which estimates them precisely from a large sample. Since datasets from other laboratories may differ from the reference in overall level and spread, each is aligned before scoring. For a cell *i* with value *yᵢ*, the reference gives the expected value *μᵢ* at that cell’s age and depth, and the standard deviation σᵢ expected for a cell from a new subject. The location offset *c* is the maximum-likelihood intercept of a linear mixed model (LMM) of the deviations *yᵢ − μᵢ*, with a random intercept for each subject, so that more heavily sampled subjects do not dominate the offset. Spread is measured with the interquartile range to be insensitive to outliers in small datasets. The scale factor is *s = IQR((yᵢ − μᵢ − c)/σᵢ)/Qref*. *Qref* is the interquartile range the reference itself shows at the same ages and depths, obtained by simulation from the fitted model. Each cell is scored as *zᵢ = (yᵢ − μᵢ − c)/(σᵢs).* Its centile is the reference’s Student-t residual distribution evaluated at *zᵢ*. The offset is tested with a Wald test and the scale factor with a subject bootstrap. Group differences are tested with an LMM of the aligned scores, with a random intercept for each subject. A remaining trend with age, tested in the same way, indicates a trajectory that differs from the reference. In a case-control design, only the offset is estimated, from control cells, and applied to all (Fig. S17b). The SHERPA resource provides our human frontal cortex dataset as a worked example (Fig. S17a), together with mouse developmental datasets from frontal^59^, auditory^60^ and visual cortex^28^ (Table S2).

---

**Figure S1.**
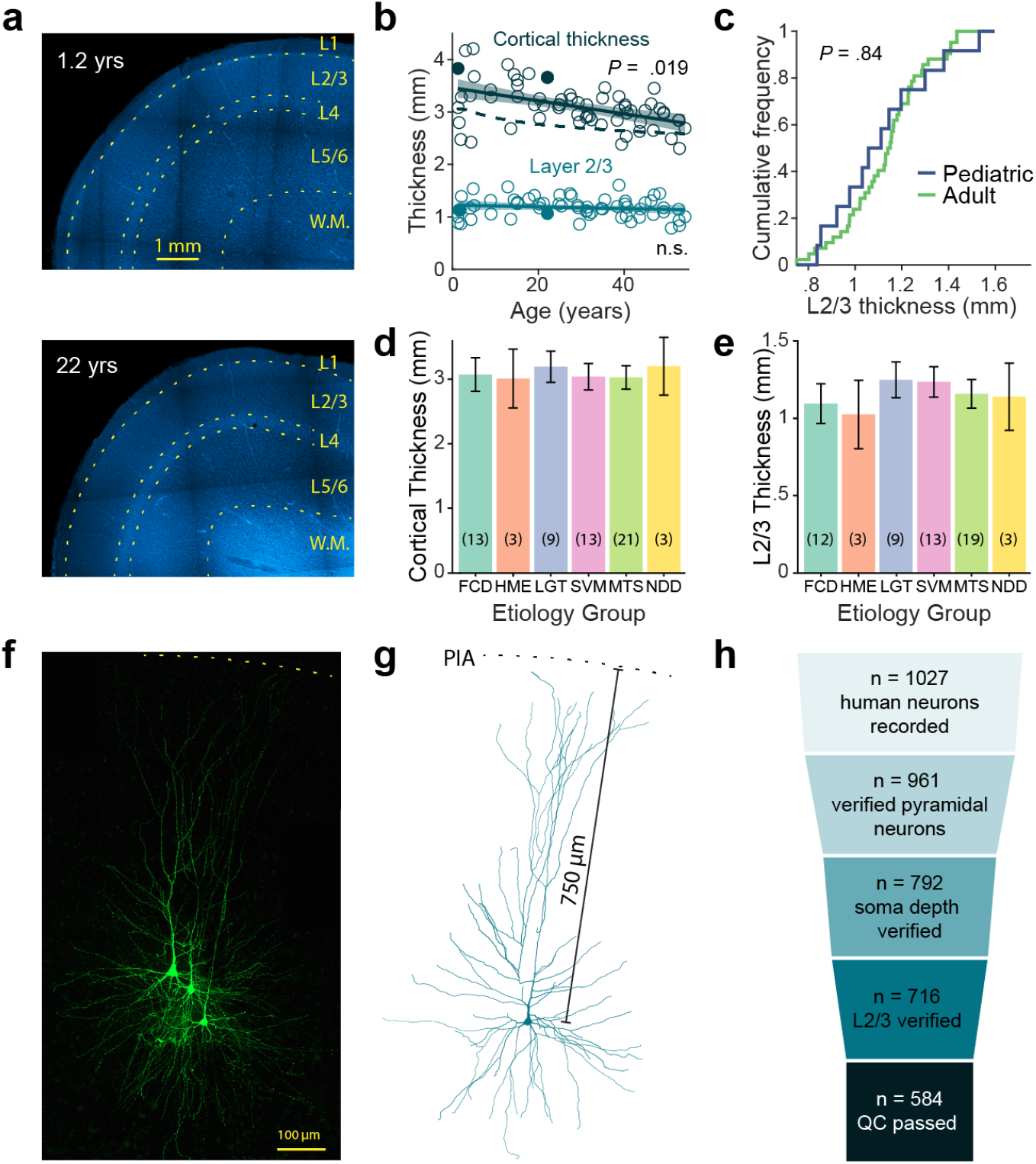
Histology and cell selection. **a**, Example images of DAPI stained cortical slices, used to estimate layer boundaries (yellow dashed) and cortical thickness. Note fainter white matter in the infant due to lower degree of myelination. **b**, Cortical thickness and L2/3 thickness estimates vs age. Cortical thickness (CT) decreases mildly with age (−12.4 µm/year; *P* < 0.001, *R²* = 0.24, 64 subjects), in line with earlier reported age-dependent cortical thinning (dashed line: MRI-based CT estimate from Bethlehem et al., 2022), in contrast to supragranular layers 2/3 (−1.6 µm/year; *P* = 0.28, *R²* = 0.003, 61 subjects). Filled circles correspond to examples in **a**. **c**, Cumulative frequency plots of pediatric (≤12 yrs) and adult (≥20 yrs) layer 2/3 thickness. No difference was observed in the distribution between age groups (Kolmogorov-Smirnov test; *P* = 0.837, *n*_ped_ = 12, *n*_adult_ = 42). **d**, Cortical thickness for different epilepsy etiologies. Values represent age-corrected group means ± 95% c.i.. No significant differences were observed between epilepsy etiologies (all pairwise Tukey-adjusted comparisons *P* > 0.84). FCD: focal cortical dysplasia, HME: hemimegalencephaly, LGT: low-grade tumors, SVM: structural/vascular malformations, MTS: mesial temporal sclerosis, NDD: neurodevelopmental disorders (Table S1). **e**, Same as **d** for supragranular layer 2-3 thickness. No significant differences were observed between epilepsy etiologies (all pairwise comparisons *P* > 0.52). **f**, Example image of neuronal morphology of recorded neurons, visualized by Alexa488-streptavidin stain of biocytin-filled neurons. **g**, 3D reconstruction of right-most cell in **f**, showing measurement of soma depth along apical dendrite. **h**, Funnel diagram presenting sequential selection criteria for neuron inclusion in the study.

**Figure S2.**
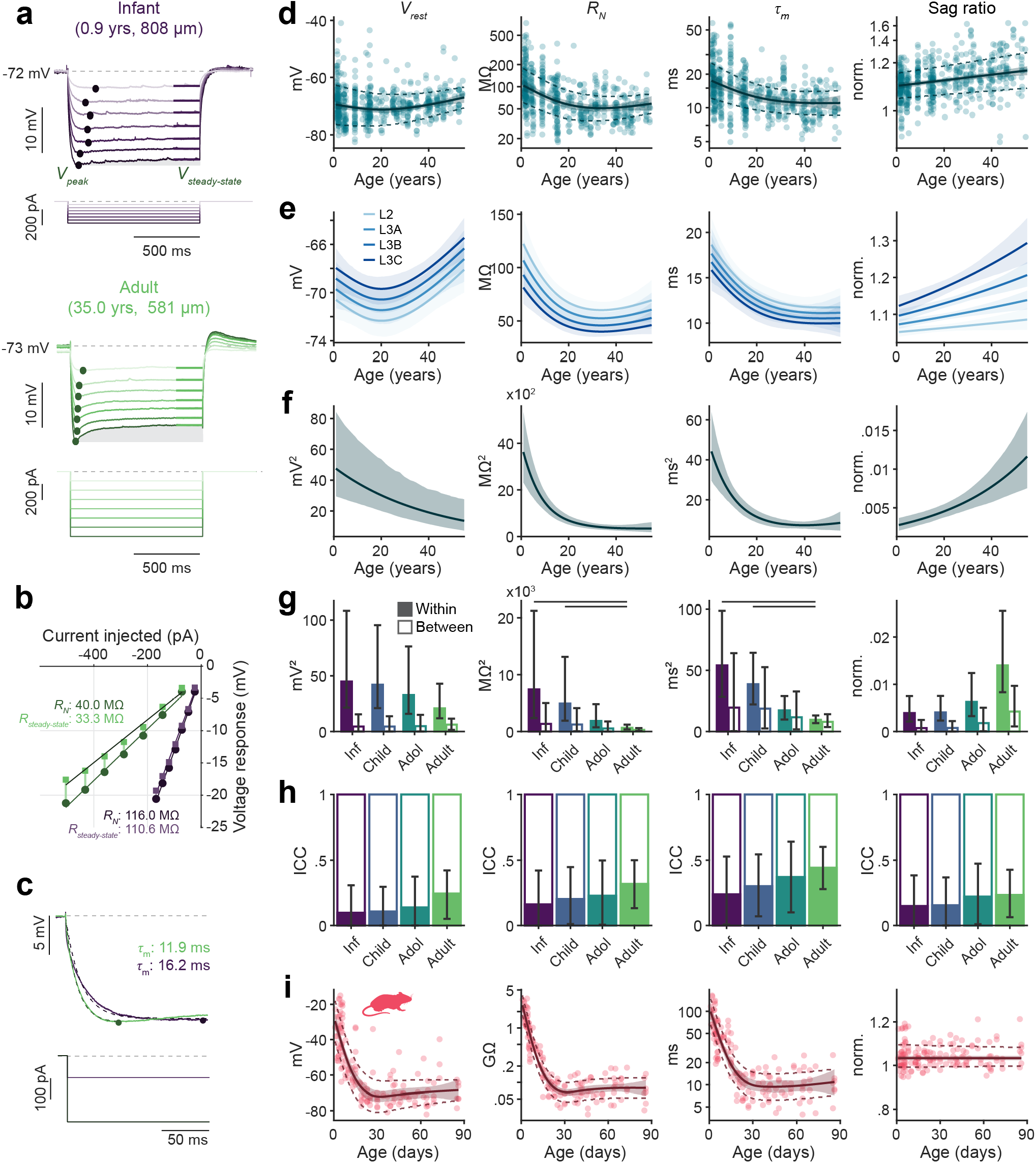
Developmental trajectories of subthreshold properties. **a**, Voltage responses (top) to hyperpolarizing current pulses (bottom) from example infant and adult neuron. Points and horizontal bars mark the peak and steady-state voltage deflections. Grey shaded area shows voltage change due to HCN-channel activation (*I*_h_), which is notably larger in the adult neurons, resulting in more prominent voltage ‘sag’. **b**, I–V relationships of the peak and steady-state deflections for both neurons. Slope of the linear fit to *V*_peak_ gives the neural resistance *R*_N_; sag ratio is defined as *R*_N_/*R*_steady-state_. **c**, Single-exponential fits from pulse onset to peak deflection, yielding the membrane time constant τ_m_. **d**, Human GAMM fits for extracted subthreshold properties, on the scale used for fitting. **e**, Trajectories for the four supragranular sublayers (L2, L3A, L3B, L3C); soma depth was a significant fixed effect in all models; for sag ratio, soma depth showed significant interaction with age (soma depth effect on sag ratio became more prominent with age). **f**, Fitted variance as a function of age, mean ± 95% c.i. from parametric bootstrap replicates. **g**, Within- and between-subject variance per age category (mean ± 95% c.i., bars indicate *P* < 0.05). **h**, Intraclass correlation coefficient (ICC) per age category (mean ± 95% c.i.). **i**, As **d**, for mouse.

**Figure S3.**
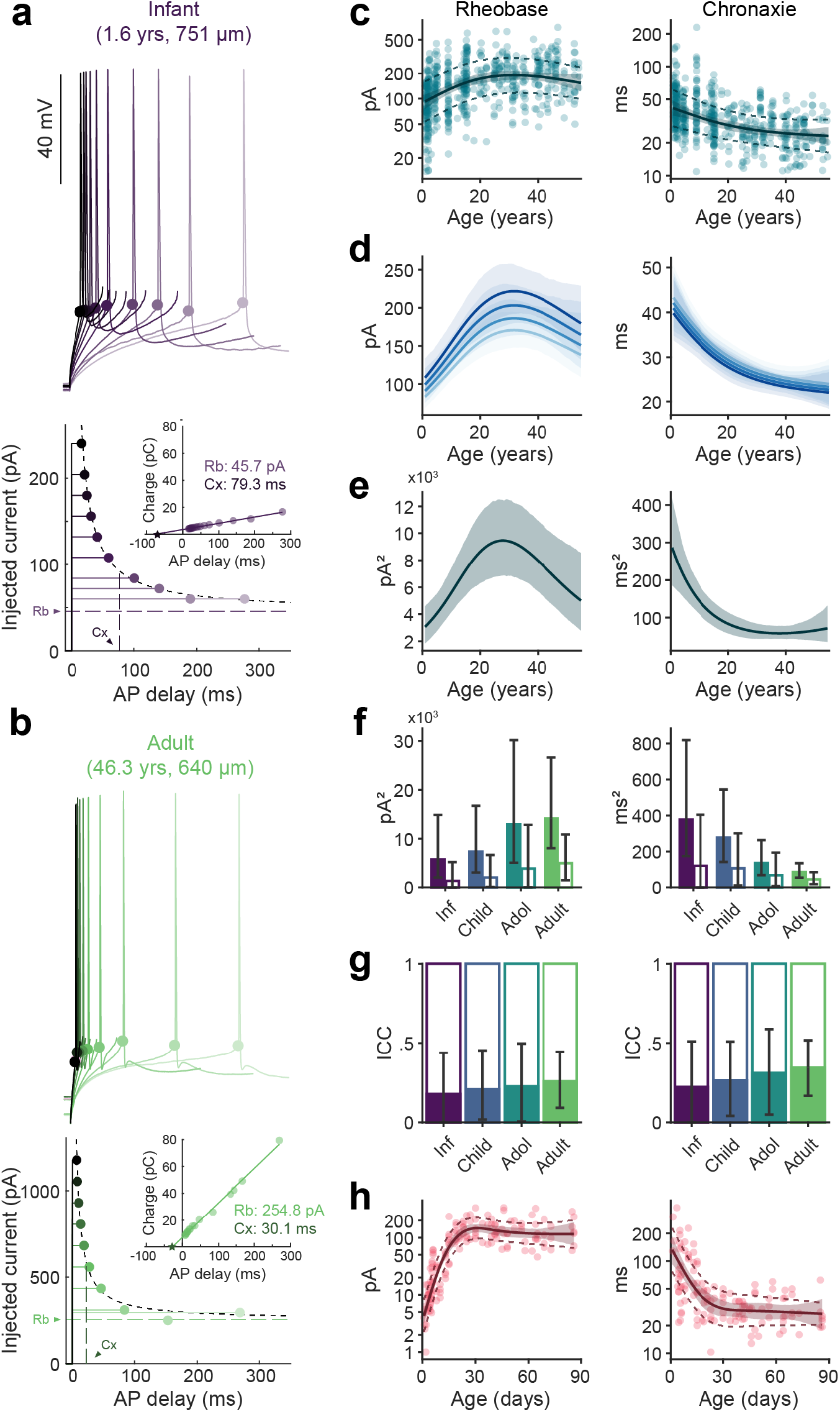
Developmental trajectories of excitability properties. **a, b**, Rheobase (Rb) and chronaxie (Cx) estimated from the relationship between injected current and delay to the first AP, for an example infant (**a**) and adult (**b**) neuron. Top: voltage responses to increasing current steps, with the first AP marked (points). Bottom: AP delay against injected current, fitted with Lapicque’s equation (dashed); inset, the same data as threshold charge against delay, from which rheobase (slope) and chronaxie (negative x-intercept, star) are read. **c**, Human GAMM fits for the excitability properties, on the scale used for fitting. **d**, Trajectories for the four supragranular sublayers (L2, L3A, L3B, L3C); soma depth was a significant fixed effect for rheobase. **e**, Fitted variance as a function of age, mean ± 95% c.i. from parametric bootstrap replicates. **f**, Within- and between-subject variance per age category (mean ± 95% c.i.). **g**, Intraclass correlation coefficient (ICC) per age category (mean ± 95% c.i.). **h**, As **c**, for mouse.

**Figure S4.**
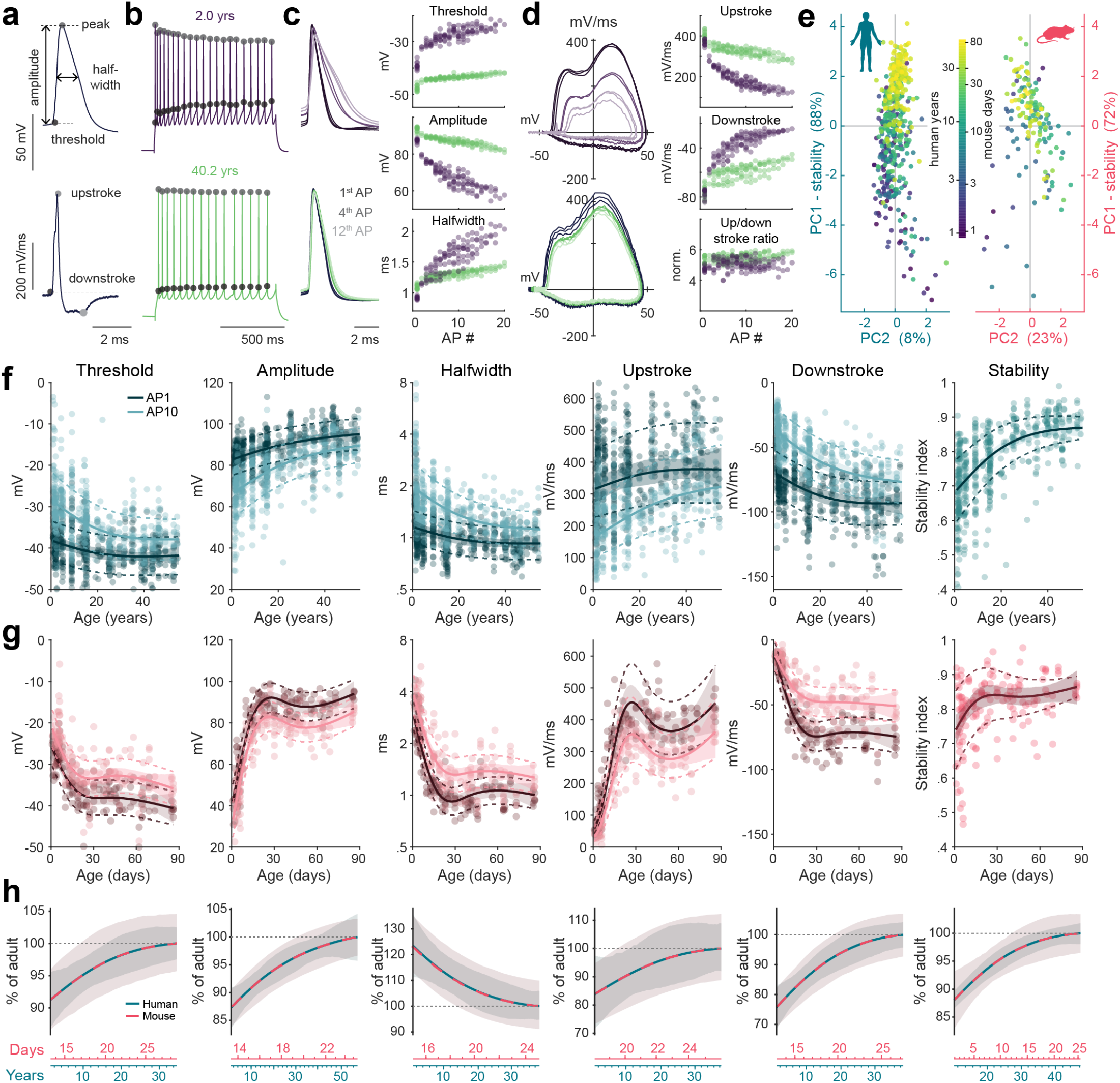
Developmental trajectories of action potential kinetics. **a**, Example AP waveform (top) and its derivative (bottom), showing the extracted kinetic features; threshold is the point where the derivative last crosses 15 mV/ms. **b**, AP trains from example infant and adult neurons, points marking threshold and peak of each AP. **c**, Left: overlay of APs at different positions in the train. Right: kinetic features of 100 APs from the same neurons, recorded across several current steps, plotted against their position in the train. **d**, As **c**, for the AP phase plane and the features of AP rise and fall. **e**, Derivation of the AP stability index: PCA over the adaptation ratios of the five kinetic features, with cells shown in PC1-PC2 space coloured by age. PC1, which accounts for 88% (human) and 72% (mouse) of the variance, is rescaled to give the index (Methods). **f**, Human GAMM fits for the AP kinetic features, shown for the first and tenth AP of the train (AP1, AP10), and for the AP stability index, on the scale used for fitting. **g**, As **f**, for mouse. **h**, Human and mouse trajectories overlaid, each expressed as a percentage of its own adult value with 95% confidence bands; the two age axes are scaled so that the curves superimpose, as in Fig. 1d.

**Figure S5.**
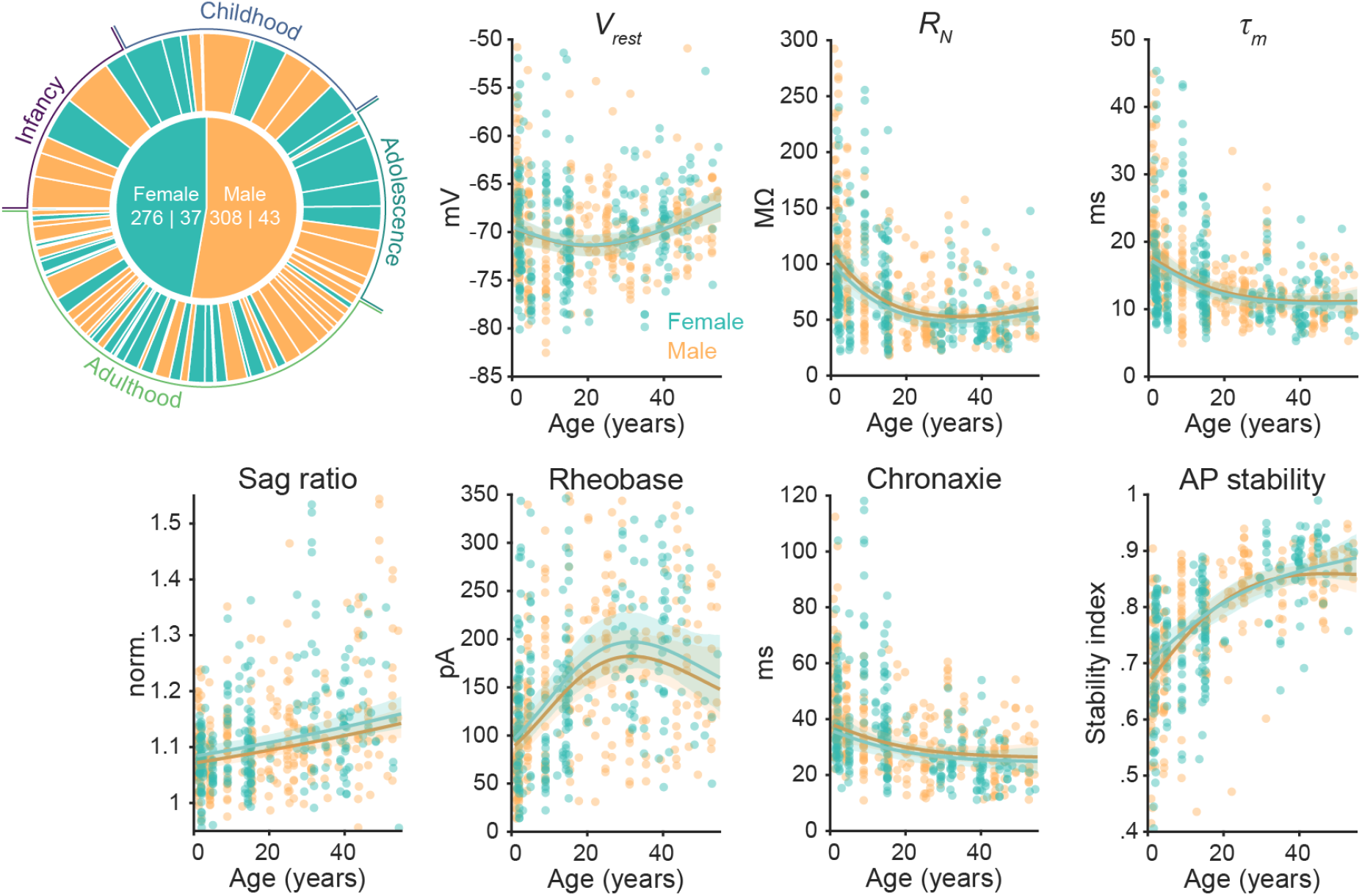
Developmental trajectories are independent of biological sex. **a**, Composition of the human dataset by sex. Inner disc: cells | subjects per sex. Outer ring: individual subjects ordered by age, with age categories indicated. **b**, GAMM fits with sex added as a fixed effect, for a representative set of features; points are individual neurons, lines with shading the predicted mean and 95% c.i. per sex. Including sex, either as an additive term or as an interaction with age, did not significantly improve model fit for any feature.

**Figure S6.**
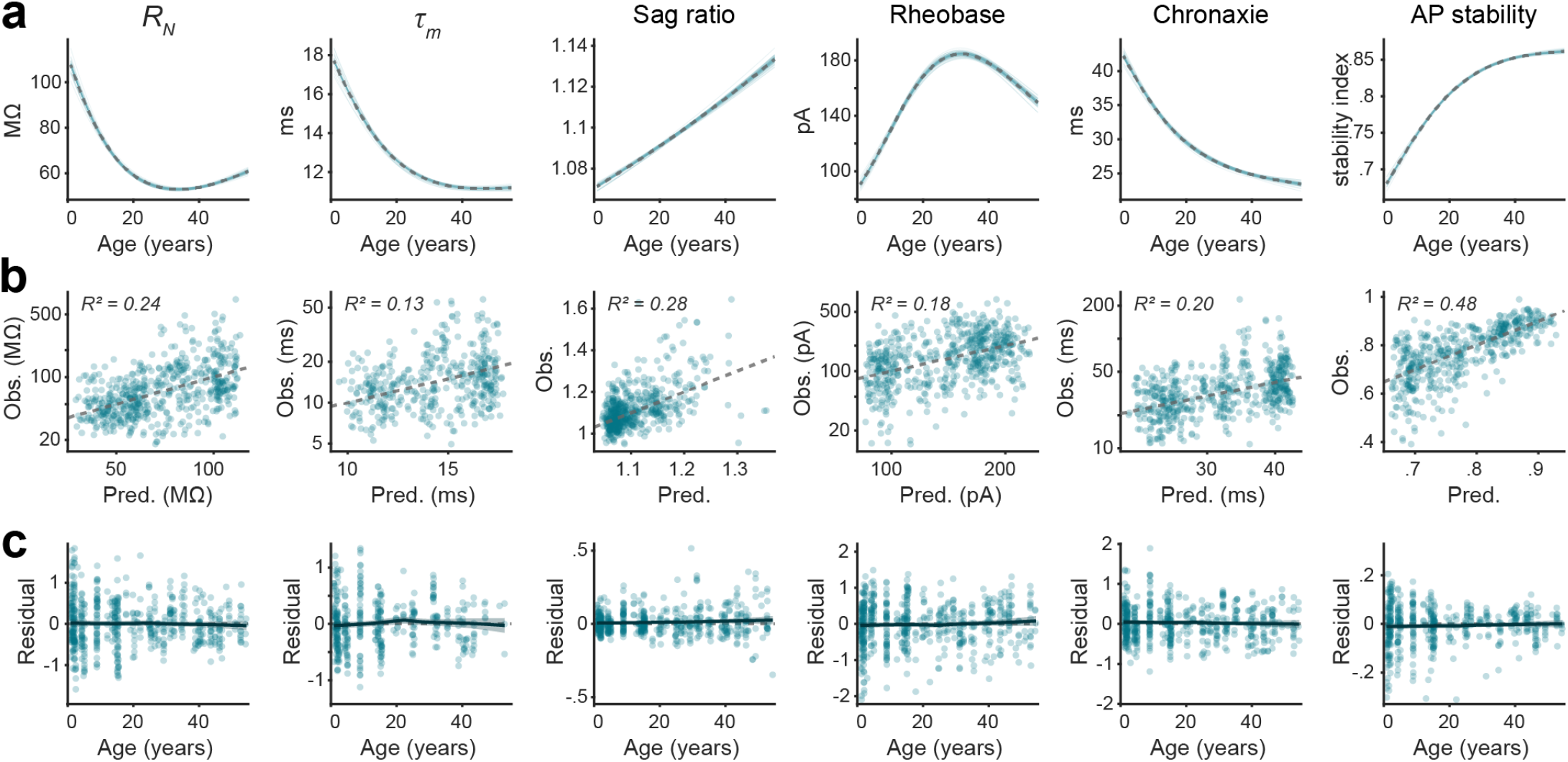
Leave-one-out validation of the human developmental models. **a**, Leave-one-subject-out refits. GAMM trajectories refitted with each subject omitted in turn (thin lines), overlaid on the full-data fit (dashed). **b**, Leave-one-out cross-validation. Observed against predicted values, on the scale used for fitting, each cell predicted from age and soma depth by a model fitted without it; dashed line, identity; R², variance explained by the cross-validated predictions. **c**, Cross-validation residuals against age, with a smooth fit and 95% c.i. showing no systematic deviation across development.

**Figure S7.**
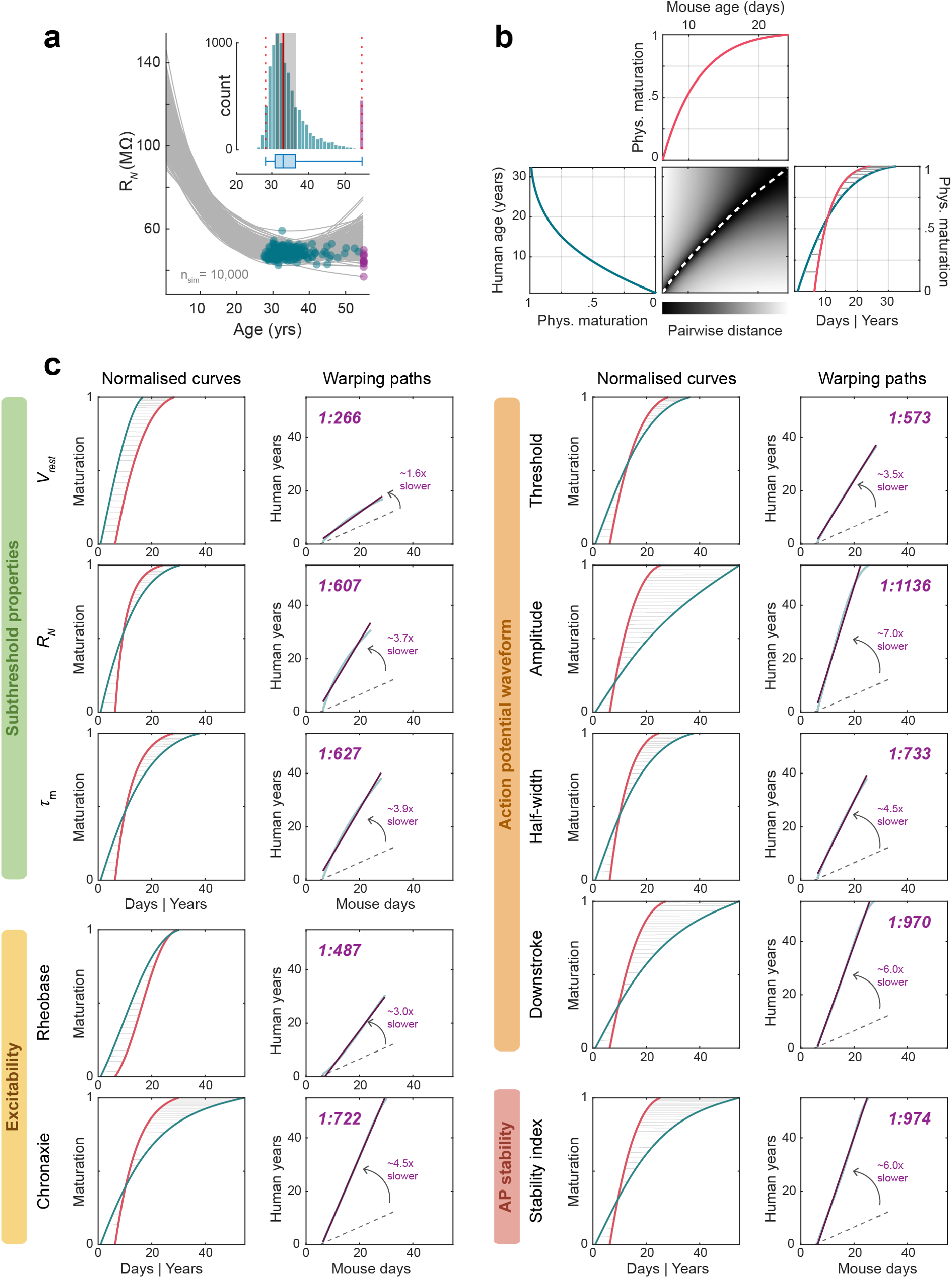
Dynamic time warping of human and mouse developmental trajectories. **a**, Peak maturation age for physiological features was estimated using 10,000 bootstrap replicate samples from the GAMM models and fit (grey lines, showing subset of 250 fits to human *R_N_* replicates), and for each the adult peak age (blue points) was determined. The distribution of peak values (inset) yielded the median peak age (red line) and provided measures of prediction accuracy in the form of IQR (shaded) and 2.5-97.5% percentiles (dashed), summarized in box plot below. Purple points mark fits where no plateau value was reached within sampled age range. **b**, Dynamic time warping, shown for *R_N_*. Human and mouse trajectories are min-max normalised over the brain growth period of each species (left, top). Heat map: pairwise distance matrix with the optimal warping path (white dashed). Right: both curves overlaid, with a subset of matched time points linked (grey). **c**, Dynamic time warping for all features with a significant age effect in both species, grouped by physiological domain. Normalised curves and warping paths with linear fits are shown as in Fig. 2c, with the scale factor, the brain growth slope (dashed) and the developmental scaling coefficient indicated for each feature.

**Figure S8.**
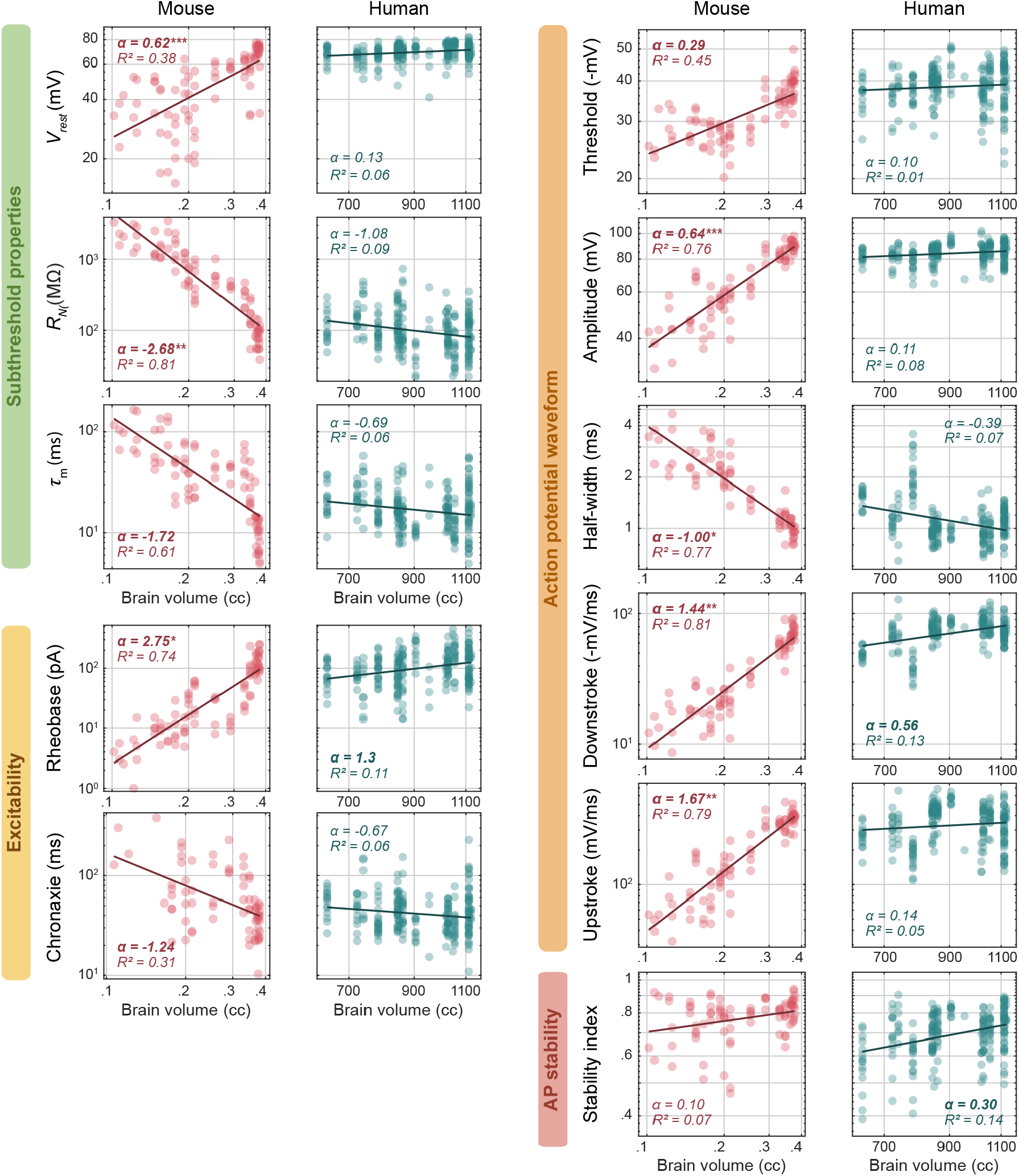
Allometric relationships between physiological features and brain volume. Allometric relationships over the brain growth phase for all features, grouped by physiological domain, as in Fig. 2e. α, allometric scaling exponent from the mixed model (Table S13), given with its marginal *R²*; bold α, 95% c.i. excludes zero; asterisks, species difference (\**P* < 0.05, \*\**P < 0.01***, \*\***\**P* < 0.001).

**Figure S9.**
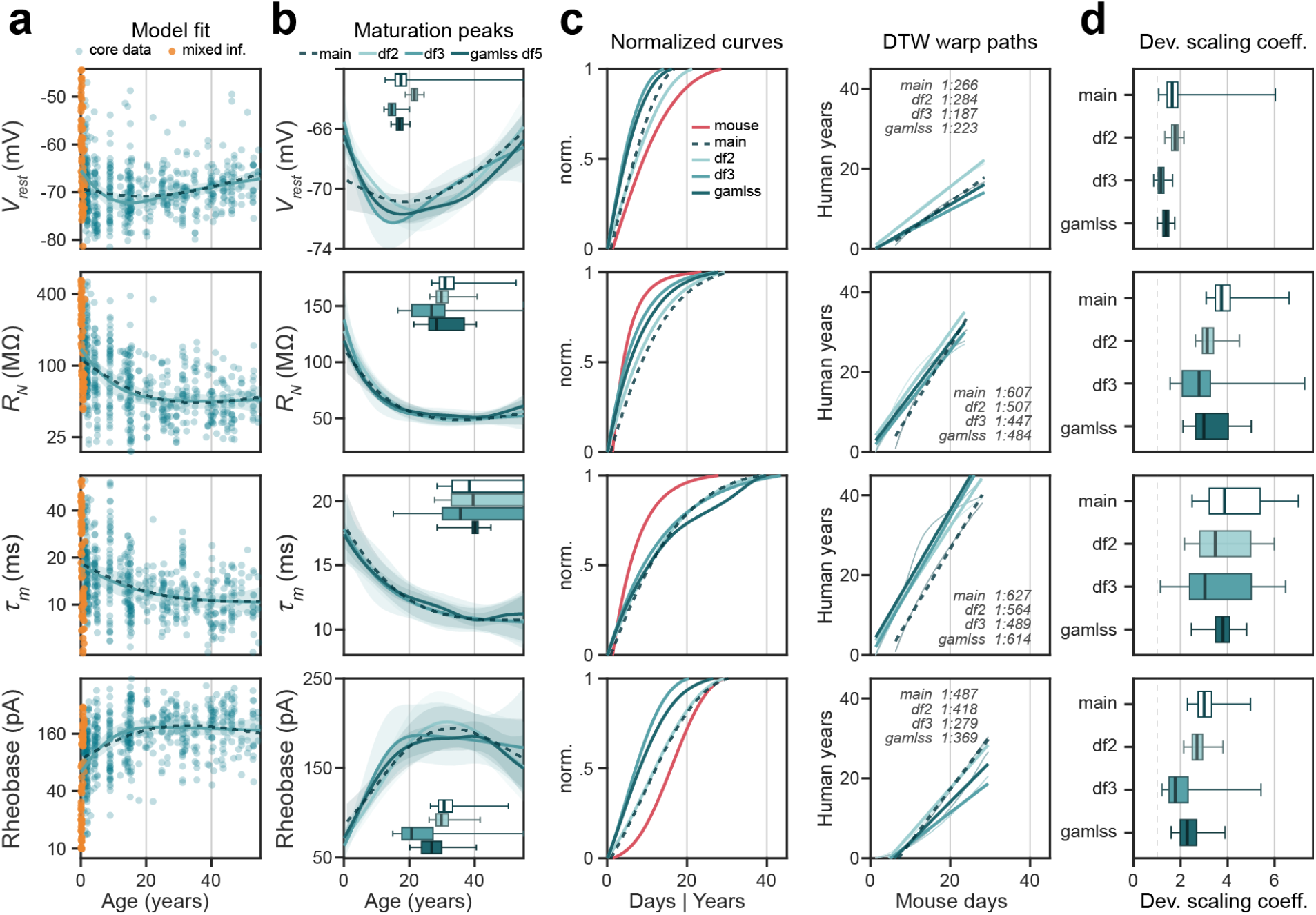
Developmental trajectories are robust to the sampled age range and model flexibility. **a**, To test whether the trajectories accurately reflected changes in early life, the core temporal cortex dataset (teal), which starts at 10 months, was extended with recently published younger infant recordings of mixed cortical origin^17^ and our own (orange; 73 cells | 9 donors, 0.08-1.29 years; Table S2), and select features were refitted. Dashed, the main model fit on core data only (dashed throughout **a** to **c**); solid, the extended data fitted with 3 df spline (mean and 95% CI). **b**, Trajectories and maturation peaks for the main model and for models fitted to the extended data with 2 or 3 spline degrees of freedom, or with a penalised spline (GAMLSS). Under model selection on the extended data the third degree of freedom was supported only for resting membrane potential (χ²(1) = 7.30, *P* = 0.007), although BIC difference was small (ΔBIC −0.8; Table S12). Median peak ages across the four specifications span 14.6 to 21.5 years for resting membrane potential, 20.7 to 30.7 for rheobase, 26.9 to 31.0 for input resistance and 35.6 to 40.2 for the membrane time constant. **c**, Normalised trajectories of each model with the mouse trajectory (red), and the corresponding DTW warping paths with linear fits; DTW scale factors indicated. **d**, Developmental scaling coefficients per model (bootstrap median, IQR and 2.5th to 97.5th percentiles). Coefficients span 1.16 to 1.77 for resting membrane potential, 1.78 to 3.01 for rheobase, 2.79 to 3.75 for input resistance and 3.04 to 3.87 for the membrane time constant. The added infant data or the additional flexibility thus did not greatly alter fitted trajectories, overall resulting in similar maturation peaks, developmental scaling coefficients, and allometric exponents (Table S14).

**Figure S10.**
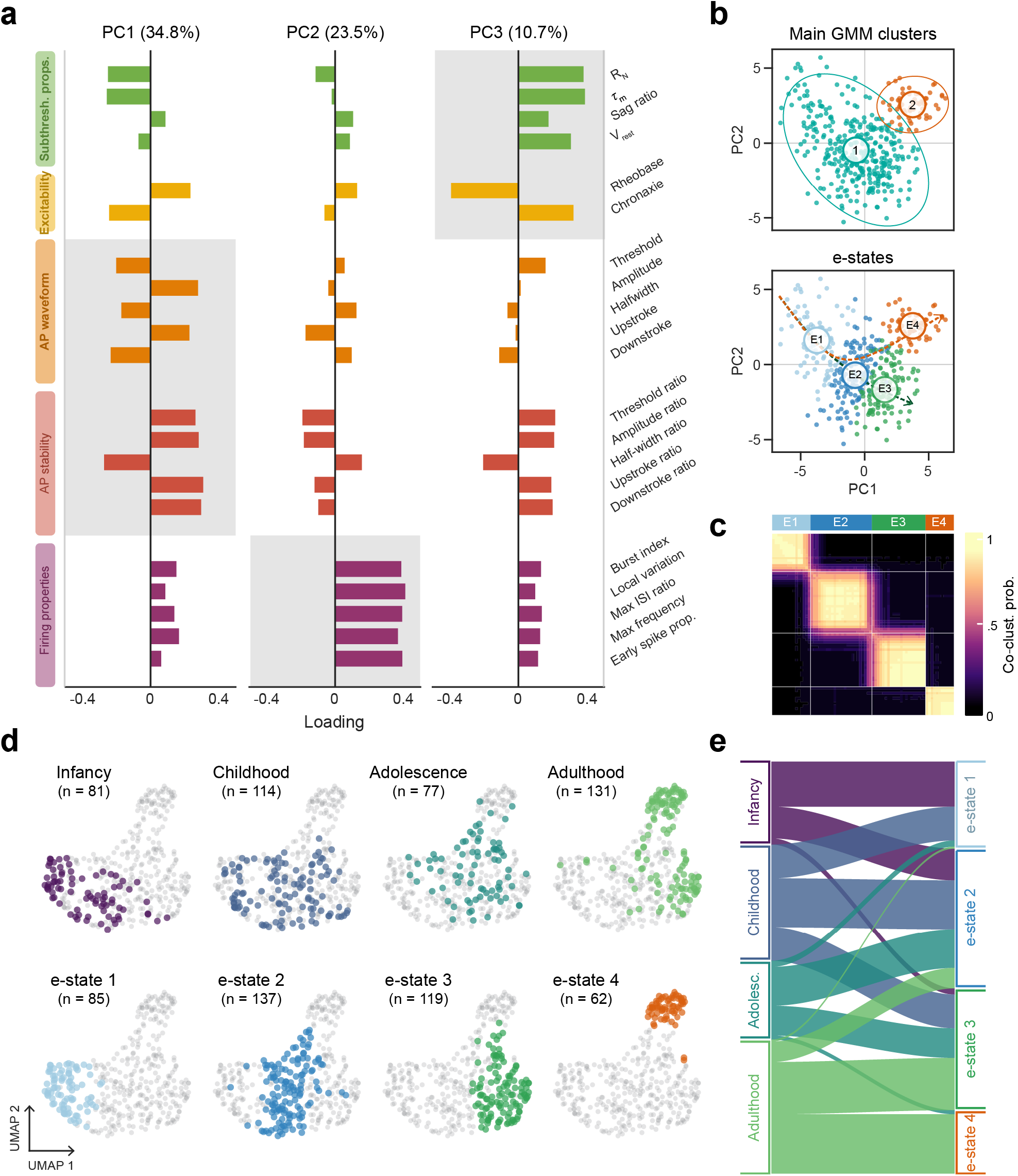
Definition of e-states along the multivariate developmental axis. **a**, PCA analysis. Loadings of the 21 features on the first three principal components grouped by physiological domain. Grey shading marks the domains loading most strongly on each component. **b**, Top: the two main clusters identified by Gaussian mixture modelling on PCA data. Cluster 1 captures a maturational axis from infancy to adulthood, cluster 2 emerges late in development (Fig. 3b). Bottom: the lineages from infancy to the two adult endpoints (dashed) and definition of e-states; main cluster 1 is subdivided by pseudotime into e-states 1 to 3 and cluster 2 forms e-state 4. **c**, Co-clustering probability of cell pairs across 1,000 bootstrap resamples, ordered by e-state. Two clusters were recovered in 96.5% of resamples, with a median Jaccard index of 0.995 for the continuum and 0.975 for the late-emerging cluster; the four e-states were reproduced with median Jaccard indices of 0.895, 0.825, 0.871 and 0.975. **d**, UMAP embedding of the 21 scaled features, with cells highlighted by age category (top) and by e-state (bottom), showing the progression from infancy to adulthood and the separation into two distinct adult e-states defined in PCA space. **e**, Alluvial plot mapping age categories to e-states; e-state 1 comprises mainly infant and childhood cells, e-state 2 draws on all age categories, and adult cells fall almost entirely in e-states 3 and 4.

**Figure S11.**
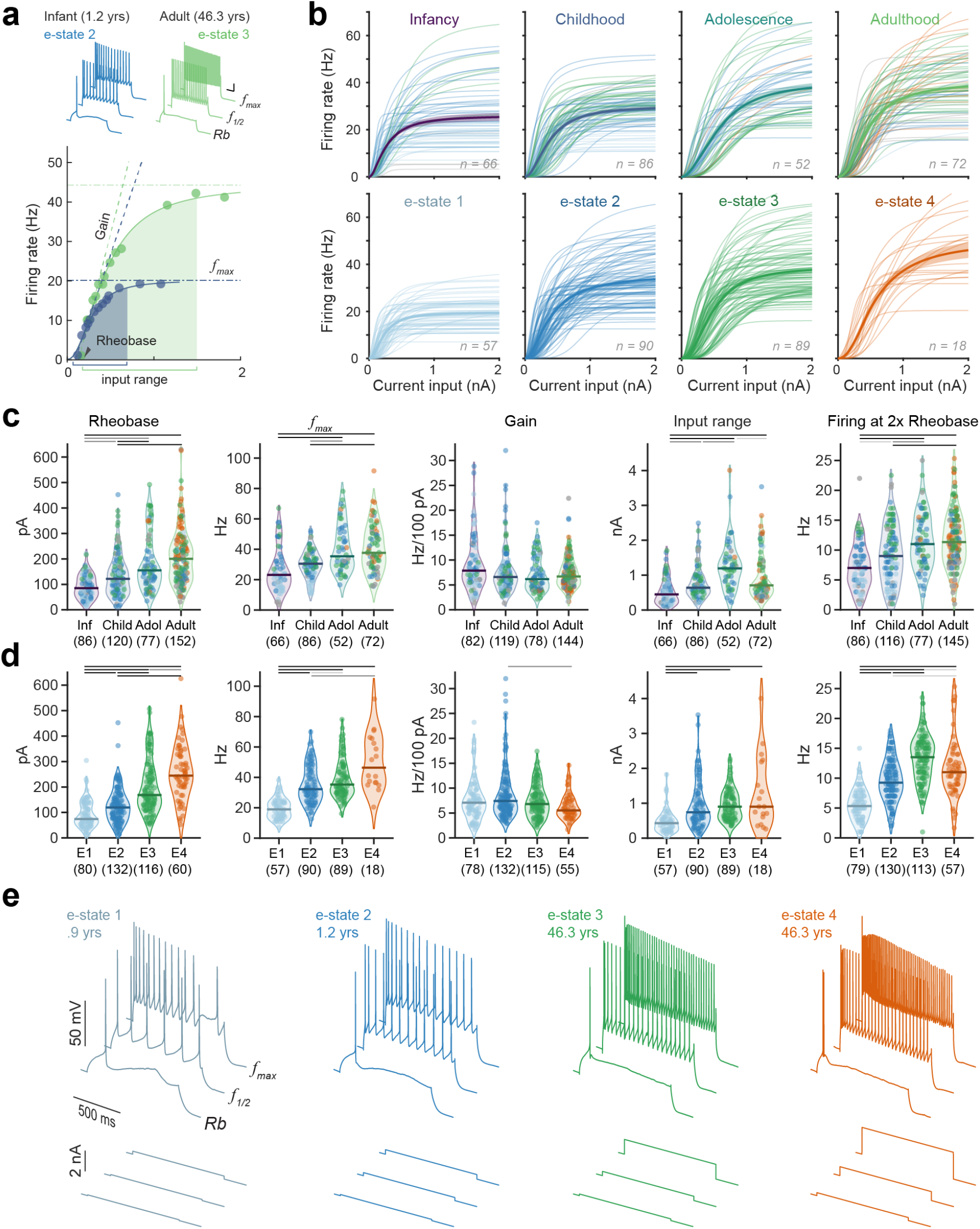
Neuronal input-output relationships change along the developmental axis. **a**, Extraction of input-output features from the firing rate against injected current (f-I) relationship, shown for an infant and an adult neuron with example traces at rheobase (Rb), half-maximal (*f*_1/2_) and maximal firing rate (*f*_max_). Curves are fits to the firing rates at each current step; gain is the initial slope (dashed), *f*_max_ the plateau (dash-dot) and input range the span of currents over which the firing rate rises (shaded). **b**, f-I curve fits of individual neurons by age category (top, coloured by e-state) and by e-state (bottom), with mean ± s.e.m. (thick). **c**, Input-output features by age category. Violins show median, *n* in brackets; top lines mark significant pairwise differences (KW with post hoc tests; black, *P* < 0.001; grey, *P* < 0.01; light grey, *P* < 0.05). **d**, as **c**, for e-states. **e**, Firing patterns at rheobase, *f*_1/2_ and *f*_max_ for one example neuron per e-state, with the corresponding current injections (bottom).

**Figure S12.**
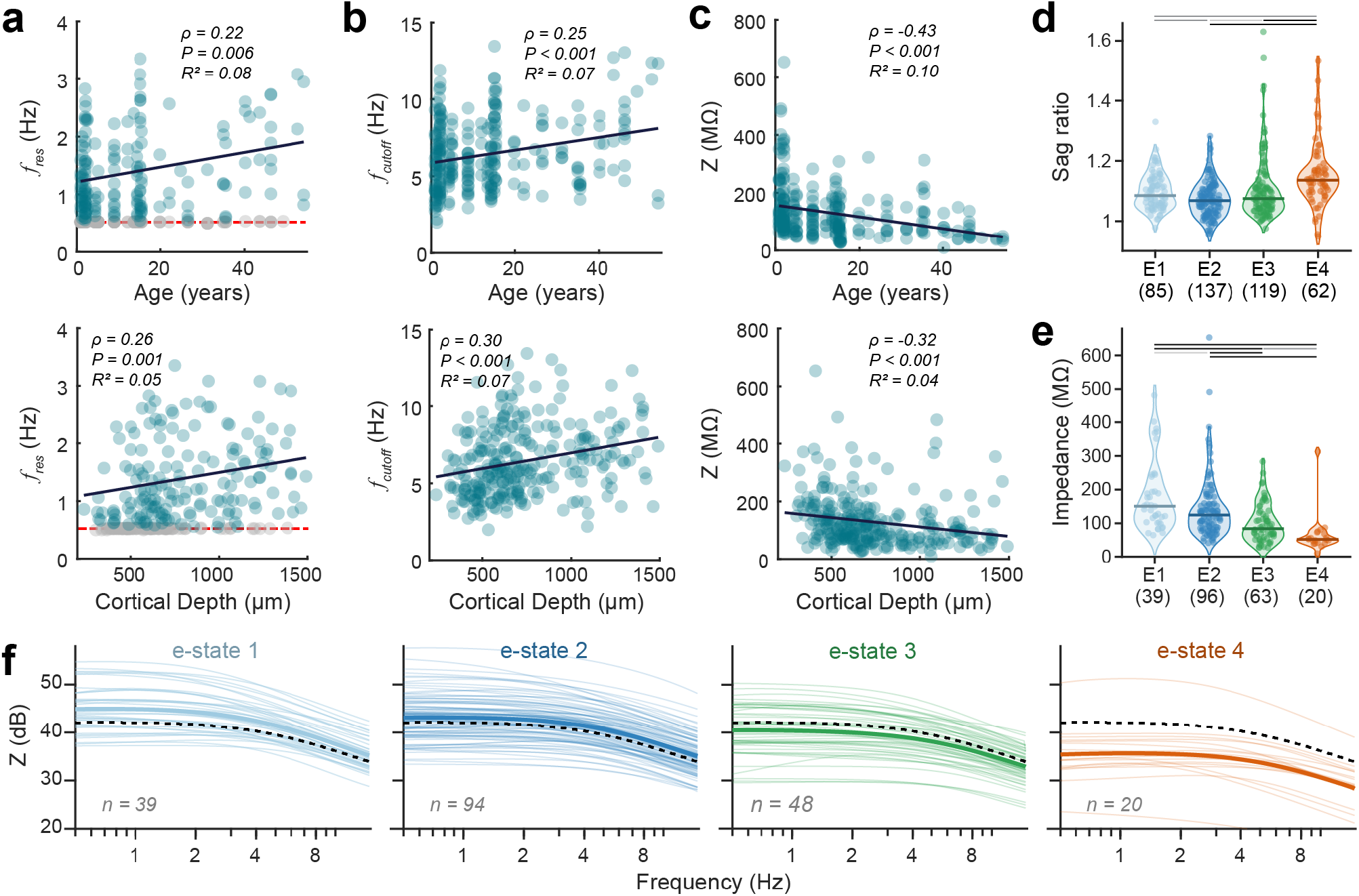
Subthreshold filtering properties mature with age and differ between e-states. **a**, Resonant frequency against age (top) and cortical depth of recording (bottom), with Spearman ρ, *P* and the *R²* of the linear fit. Grey points mark neurons with resonance at the lowest frequency tested (0.5 Hz, red dashed). **b**, As **a**, for the 3 dB cutoff frequency. **c**, As **a**, for the average impedance below the cutoff frequency. All three properties change significantly with age and soma depth. **d, e**, Sag ratio (**d**) and impedance (**e**) per e-state; violins with median, *n* in brackets; lines mark significant pairwise differences (KW with post hoc tests; black, *P* < 0.001; grey, *P* < 0.01; light grey, *P* < 0.05). E-state 4 has the highest sag ratio and the lowest impedance. **f**, Bode magnitude plots of impedance against frequency (log scale) per e-state, in dB re 1 MΩ to display the full range of response profiles; thin lines, individual neurons; thick, e-state mean; dashed, population average. Note the markedly lower impedance of e-state 4 across the sampled frequency range.

**Figure S13.**
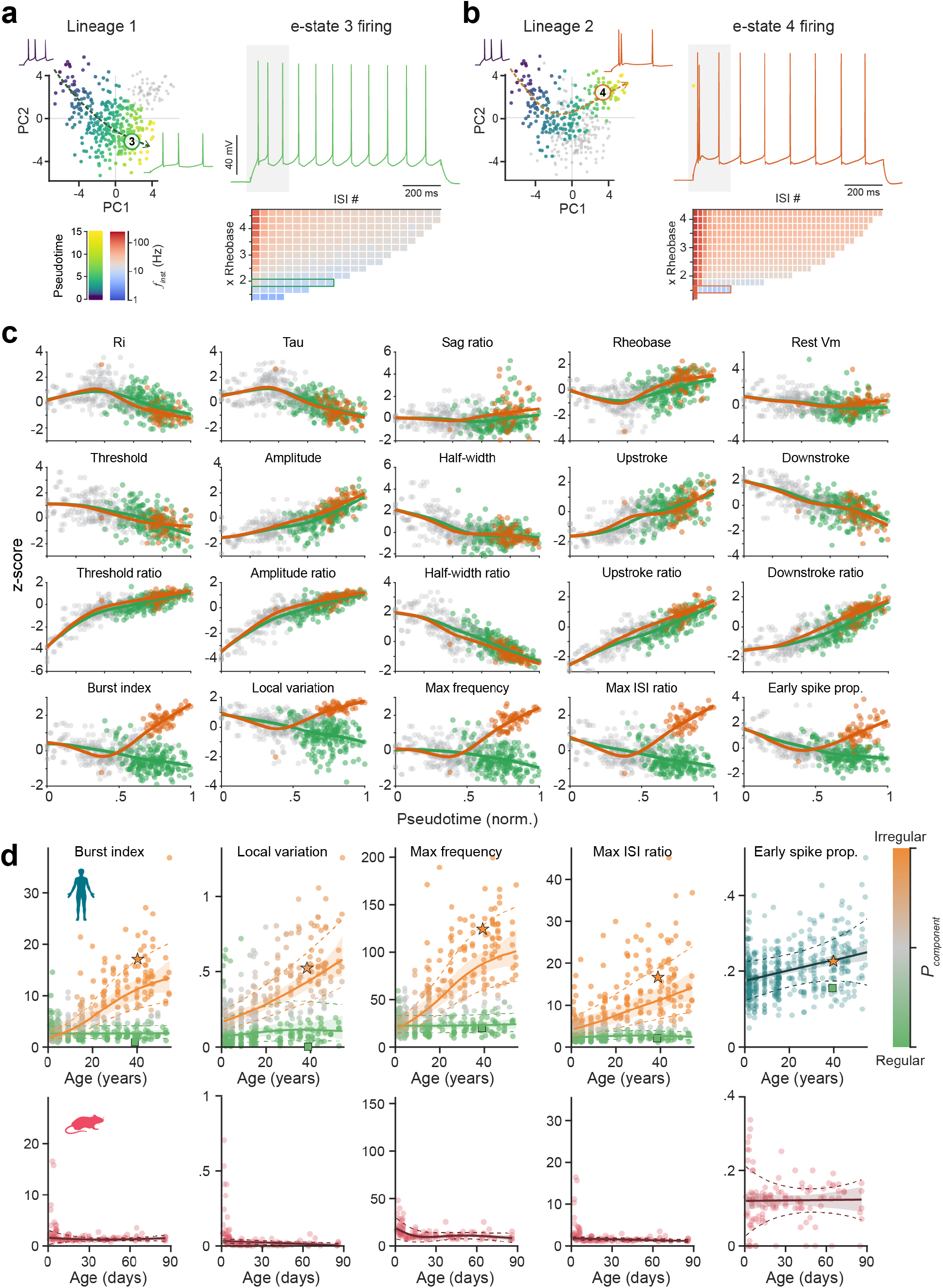
Late emergence of the adult burst-firing phenotype. **a, b**, Lineage 1 (**a**) and lineage 2 (**b**) in PC1-PC2 space, cells coloured by pseudotime (grey, not assigned), ending in e-state 3 and e-state 4 respectively, with example firing at the start and end of each lineage (insets). Right: firing behaviour of example endpoint neurons with the instantaneous firing frequency per interspike interval (ISI) across stimulus intensities below (multiples of rheobase; row of the example trace boxed). Lineage 1 ends with regular firing e-state 3 neurons, whereas lineage 2 leads to the e-state 4 neurons that initiate firing at every intensity with a >100 Hz burst. APs in grey shaded area expanded in the inset. **c**, Z-scored features against normalised pseudotime for lineage 1 (green) and lineage 2 (orange); grey, cells on the shared part of the trajectory. The lineages share similar trajectories for subthreshold, excitability, AP waveform and AP stability features but diverge markedly in firing properties (bottom row). **d**, GAMLSS mixture models of the five firing properties against age in human (top) and single-population fits in mouse (bottom). Points are coloured by the probability of belonging to the irregular (orange) or regular (green) component; lines and shading, component means with 95% c.i.; dashed, ± 1 s.d. Human early spike proportion was best described by a single component (teal, Table S7-8). Outlined star and square mark the example neurons of **b** (e-state 4) and **a** (e-state 3). The irregular, bursting phenotype appears absent in mouse supragranular layers.

**Figure S14.**
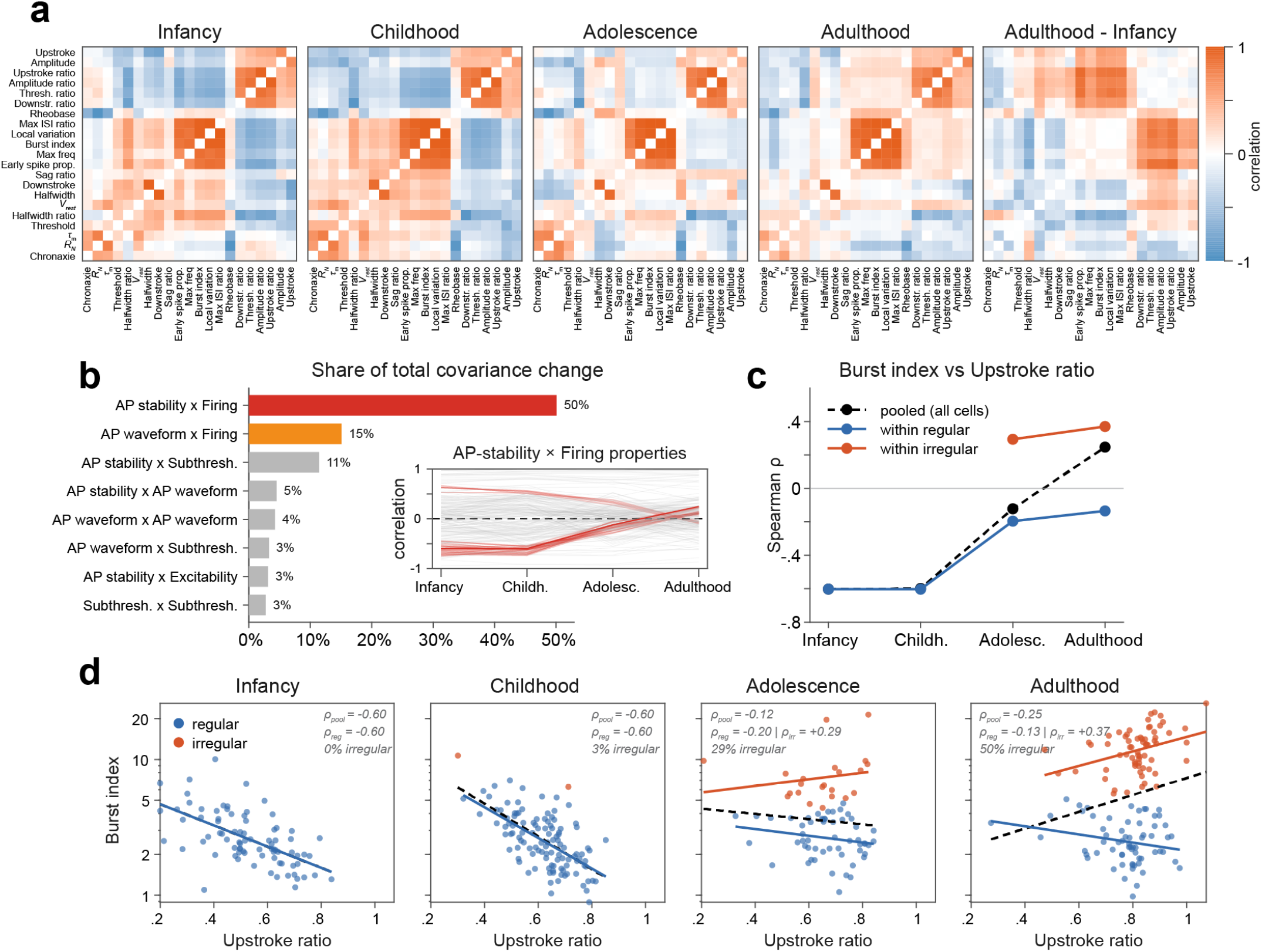
The correlation structure between physiological features reorganises with age. **a**, Pairwise Spearman correlations between the 21 features within each age category, and the difference between adulthood and infancy (right). The largest changes are between the AP stability and firing properties. **b**, Share of the total change in correlation between infancy and adulthood contributed by each pair of physiological domains. For each feature pair, the change was taken as the linear trend of its correlation across the four age categories; the share of a domain pair is the sum of squared changes of its feature pairs over the total. Inset: correlation of each AP stability and firing property pair across age categories (red; grey, all other pairs), which reverses from negative in infancy to weakly positive in adulthood. **c**, Burst index against AP upstroke ratio as an example: Spearman ρ per age category for all cells pooled (dashed) and within the regular (blue) and irregular (orange) components of the mixture models (Fig. 3g, Fig. S13d). **d**, Data underlying correlations in **c** per age category. The reversal of the pooled correlation reflects the emergence of the irregular component, in which the two features are positively correlated.

**Figure S15.**
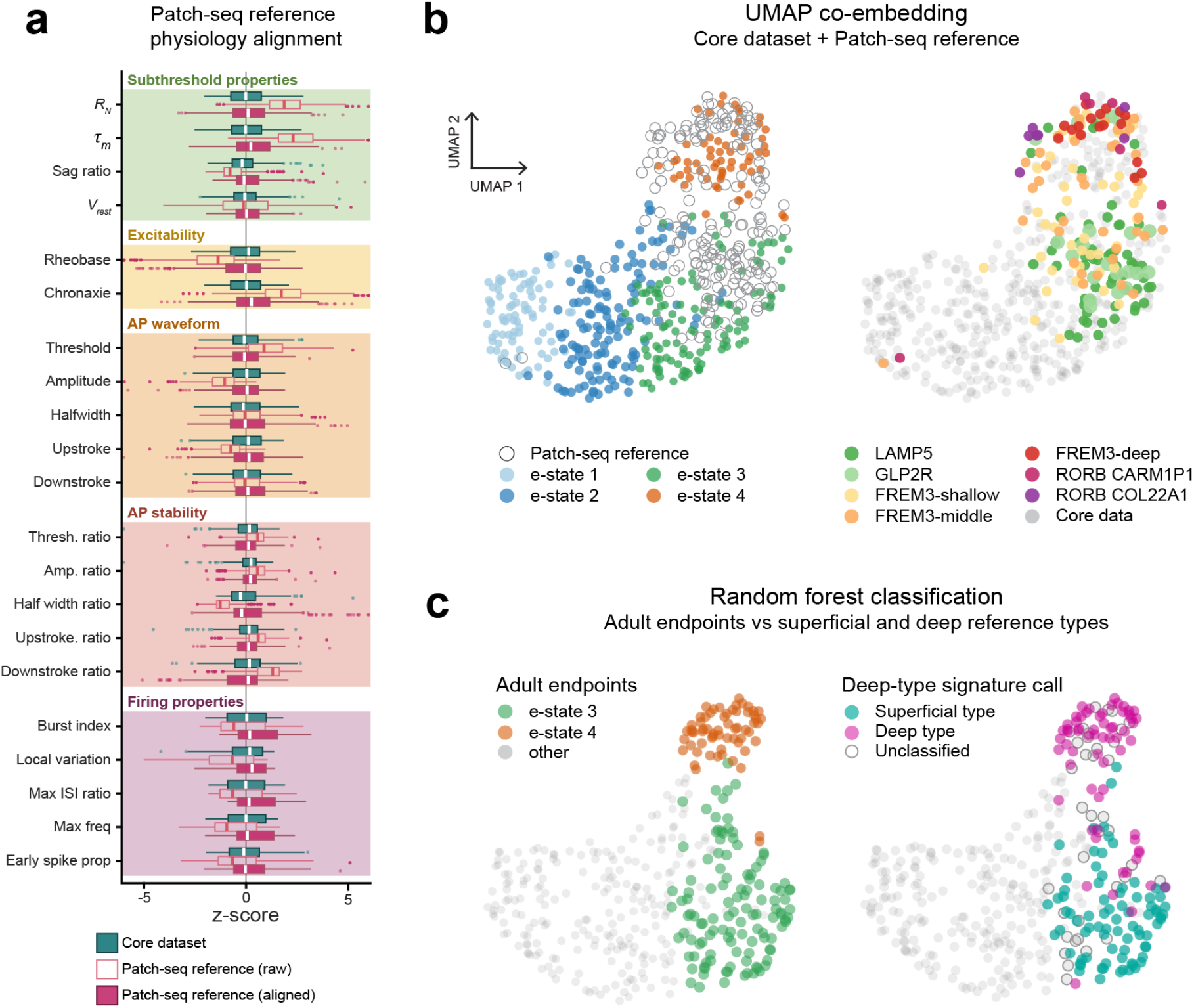
The late-emerging e-state carries the physiological signature of deep-layer transcriptomic types. **a**, Alignment of the adult human Patch-seq reference dataset (Berg et al., 2021) to the core dataset. For each of the 21 features, reference values were rescaled on the modelling scale so that their median and IQR match those of the core adult cells; box plots show the core adult cells and the reference before and after alignment, in z-scores of the core adult distribution. The aligned data were used for the projection into PCA space (Fig. 3j). **b**, UMAP co-embedding of the core dataset and the aligned reference neurons, with core cells coloured by e-state (left) and reference neurons by t-type (right). Reference neurons intermix with the core adult cells and spread over both adult endpoints; the deep types (FREM3-deep, RORB CARM1P1, RORB COL22A1) land predominantly in the e-state 4 region. **c**, Left: the adult endpoints e-state 3 and e-state 4 in the same embedding. Right: random forest classification of the e-state 3 and 4 cells, using a classifier trained on the LAMP5 and RORB reference types (Fig. 3k). E-state 4 is classified almost entirely as deep-type physiology.

**Figure S16.**
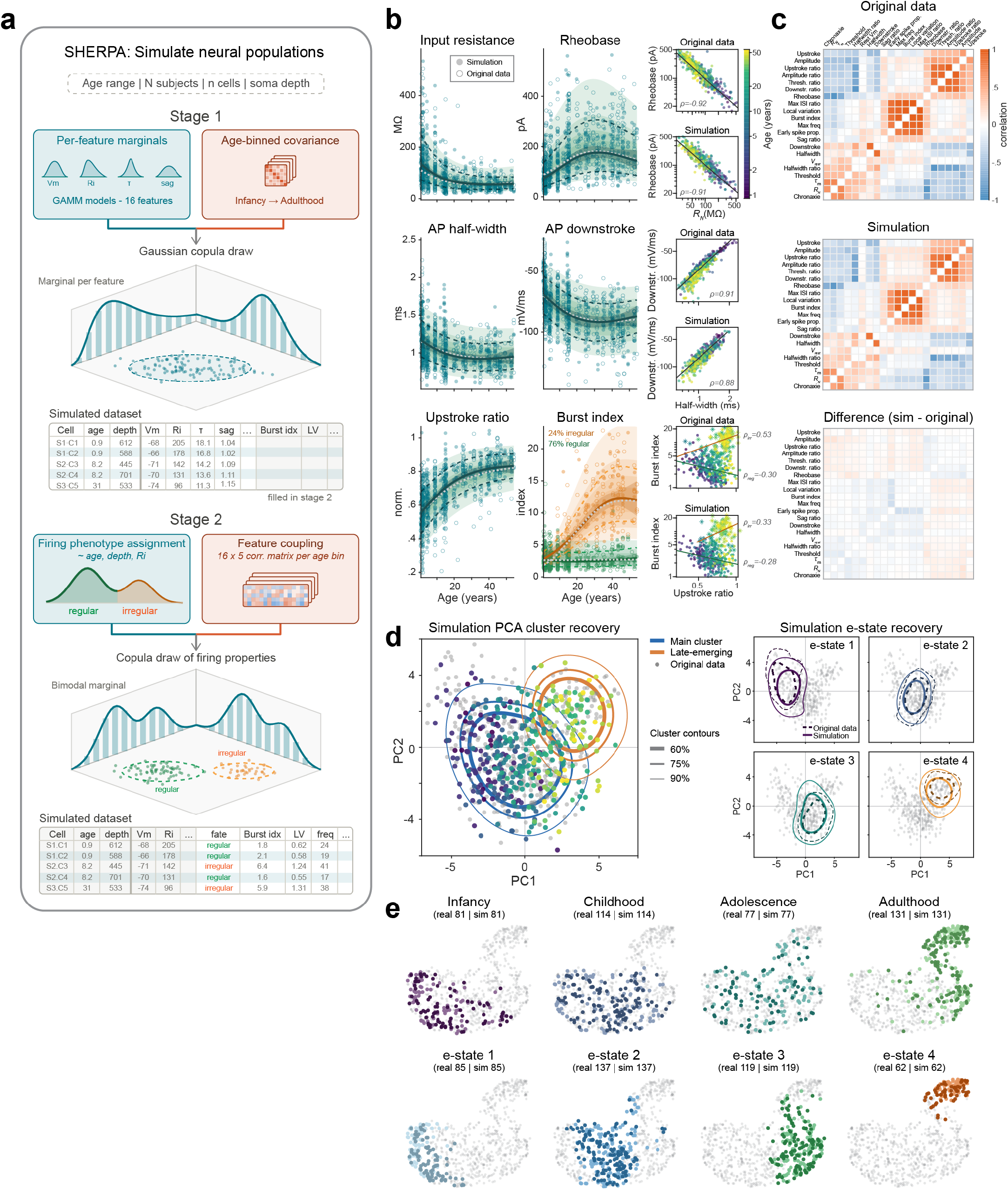
SHERPA population simulation reproduces the structure of the recorded population. **a**, The 21 GAMMs describing the physiological maturation trajectories of individual features were unified into a single generative model of the human supragranular population, from which neural populations of any age range, soma depth and size can be generated. The schematic shows the two-staged procedure in humans: for each cell, 16 features are drawn together, carrying the correlation structure of the cell’s age category; the cell is then assigned a regular or an irregular firing phenotype, and its five firing properties are drawn conditional on that assignment and on the features already drawn (Methods). **b**, A representative simulated population (filled), matched to the recorded dataset (open) in its sampling structure. Solid lines, GAMM fit to the simulated data; dashed, ± 1 s.d.; grey dashed, the reference trajectory; shading, reference centile bands. Right: pairwise feature relationships in the recorded and simulated data with Spearman ρ, showing that correlations within and between domains are preserved. **c**, Feature correlation matrices of the recorded data and of the simulation (mean over 1,000 populations matched to the core dataset in sampling structure), and their difference. **d**, Simulated cells in the PCA space of the recorded data (grey), with contours (60, 75 and 90%) of the main and late-emerging clusters; right, contours per e-state for recorded (dashed) and simulated (solid) cells. Across 1,000 simulations passed through the clustering pipeline, two clusters were recovered in 95.3% and the split reproduced the original in 82.3% (median Jaccard 0.760 for the late-emerging cluster), which comprised a median 16.9% of cells against 15.4% in the recorded data; correlation matrices matched with r = 0.97 overall and 0.83 to 0.91 within age categories. **e**, UMAP co-embedding of original dataset with matched simulation; simulated cells dark, recorded cells light.

**Figure S17.**
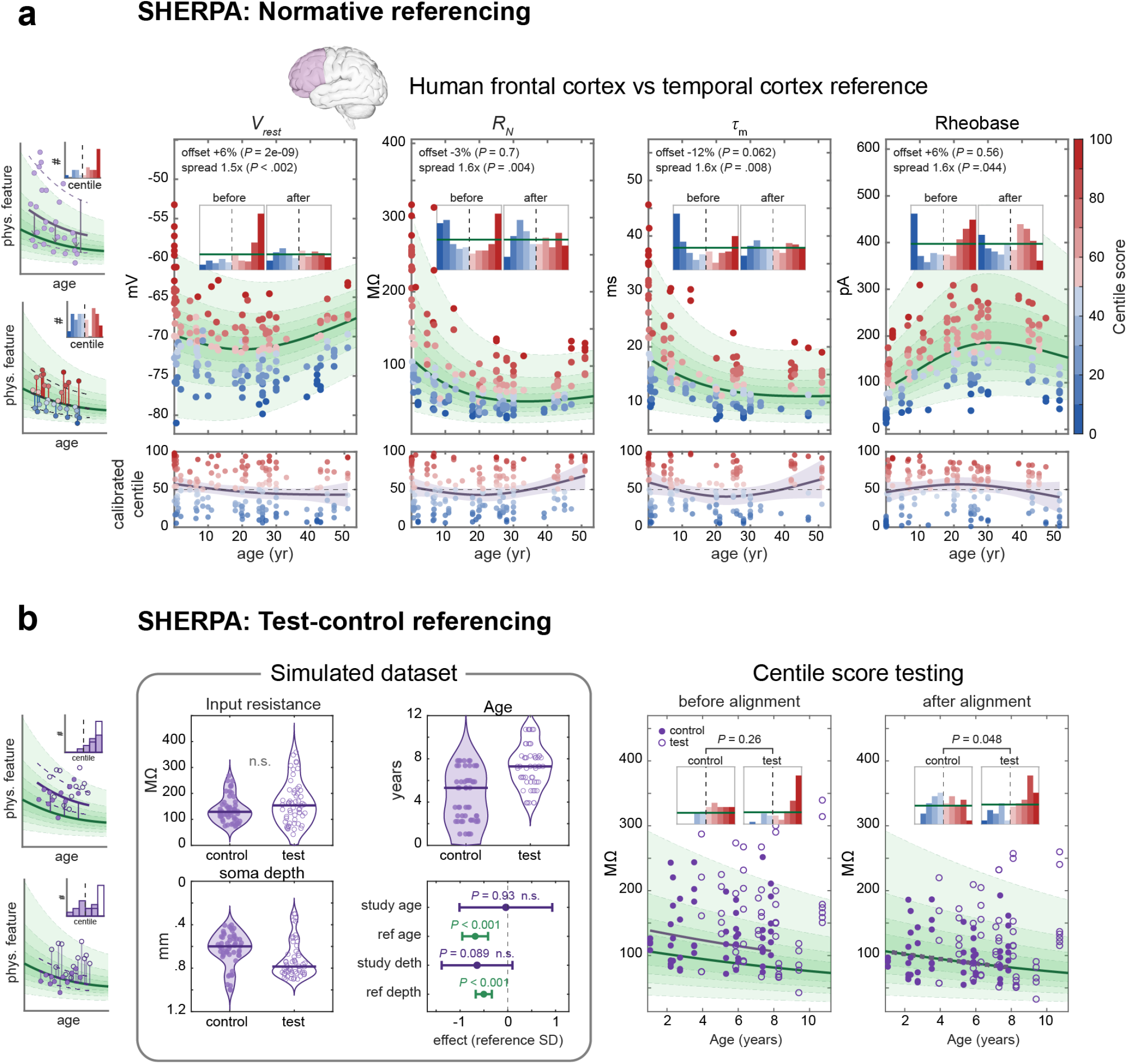
Normative and test-control referencing with SHERPA reference charts. External smaller datasets can be referenced against the GAMMs presented here in the manner of a growth chart. Each cell is scored as a centile, with the effects of age and soma depth and the expected variance taken from the reference, which estimates them more precisely from its larger sample. Because an external dataset may also differ from the reference in location and scale, it is aligned before scoring. Left schematic: A study dataset is aligned to the reference by a location offset, estimated by maximum likelihood, and a spread factor, and each neuron is then assigned a centile within the reference population at the same age and soma depth; a departure from the reference trajectory appears as an age-dependent trend in the centiles. **a**, Normative referencing example of a human frontal cortex dataset (192 cells from 44 subjects, 0.1 to 51 years; combining cells from Barzó et al., the Allen Cell Types Database, and own recordings, Table S2) against the temporal cortex reference. Main panels: frontal cells coloured by centile against the reference trajectory and centile bands for four features, with the location offset and spread factor and their *P* values; insets, centile histograms before and after alignment (green line, expected uniform distribution). Below: centiles against age with fit and 95% c.i.. No significant age-dependent trend is present (*P* = 0.09-0.30), indicating that the frontal trajectory resembles that of temporal cortex for these features. **b**, Many applications compare a test group against controls, carriers of a somatic variant against non-carriers, or cells stratified by degree of pathology. Where a control group is available it can serve as the calibration subset from which the offset is estimated, and that alignment is then applied to the whole dataset, so that control and test cells alike receive centiles on a common scale whose distributions can be compared directly. Here, test-control referencing is demonstrated using a simulated dataset in which the test group differs from controls in input resistance but is also older and sampled deeper. Within the study the group difference is not detectable (n.s.), and the effects of age and depth cannot be resolved from its own cells (study age, study depth), whereas the reference resolves both (ref age, ref depth; effects in reference s.d.). Right: centile distributions of control and test cells before and after alignment, with the control group as calibration subset; the difference becomes detectable after referencing (*P* = 0.047).

## Supplementary Tables

**Table S1.** Donor sample. Neural recordings (N = 584) were obtained from 80 tissue donors with diverse epilepsy etiologies. The sample included mesial temporal sclerosis (MTS, n = 24), structural/vascular malformations (SVM, n = 17; including cavernoma and encephalocele), focal cortical dysplasia (FCD, n = 14), hemimegalencephaly (HME, n = 4), low-grade tumors (LGT, n = 10; including ganglioglioma and DNET), and neurodevelopmental disorders (NDD, n = 5; Rasmussen’s encephalitis and Sturge-Weber syndrome). Etiology was unknown in 1 case (non-lesional) and unavailable for 5 donors. Sex is biological sex (F, female; M, male); hemisphere is given as L (left) or R (right).

| Donor # | Sex | Age (yrs) | Hemisphere | Epilepsy yrs | Etiology | # Cells | Donor # | Sex | Age (yrs) | Hemisphere | Epilepsy yrs | Etiology | # Cells |
| --- | --- | --- | --- | --- | --- | --- | --- | --- | --- | --- | --- | --- | --- |
| 1 | M | 0.9 | L | 0.9 | HME | 17 | 41 | F | 29.3 | L | 16.6 | MTS | 8 |
| 2 | M | 1.2 | R | 1.2 | HME | 13 | 42 | F | 29.6 | L | 27.6 | MTS | 2 |
| 3 | M | 1.3 | L | 1.3 | FCD | 9 | 43 | M | 31.2 | R | 11.2 | FCD | 11 |
| 4 | F | 1.6 | L | 1.4 | FCD | 23 | 44 | F | 31.3 | R | 10.1 | MTS | 6 |
| 5 | M | 2.0 | R | 1.9 | FCD | 27 | 45 | F | 31.4 | L | 5.3 | SVM | 1 |
| 6 | F | 2.0 | L |  | LGT | 12 | 46 | F | 31.9 | R | 25.6 | SVM | 5 |
| 7 | F | 2.1 | L | 2.1 | HME | 21 | 47 | F | 32.4 | L | 17.8 | MTS | 9 |
| 8 | F | 2.5 | R | 2.3 | NA | 10 | 48 | M | 34.7 | L | 12.3 | MTS | 4 |
| 9 | F | 3.0 | L | 3.0 | FCD | 4 | 49 | F | 35.0 | R | 16.0 | SVM | 6 |
| 10 | M | 4.1 | L | 3.3 | FCD | 7 | 50 | M | 35.5 | L | 12.5 | LGT | 8 |
| 11 | M | 4.3 | R | 2.2 | SVM | 1 | 51 | M | 38.4 | R | 10.3 | MTS | 1 |
| 12 | M | 4.3 | R | 1.6 | FCD | 26 | 52 | F | 38.9 | L | 32.3 | MTS | 7 |
| 13 | F | 4.8 | L | 2.4 | NDD | 2 | 53 | M | 39.4 | R | 6.3 | MTS | 2 |
| 14 | F | 5.1 | R | 4.6 | FCD | 18 | 54 | F | 39.8 | R | 24.0 | MTS | 8 |
| 15 | M | 8.9 | R |  | NA | 16 | 55 | F | 40.2 | L | 18.3 | NA | 5 |
| 16 | M | 8.9 | L | 1.7 | NDD | 14 | 56 | F | 40.2 | R | 16.7 | MTS | 1 |
| 17 | F | 9.0 | R | 5.0 | NDD | 18 | 57 | F | 40.3 | R | 6.4 | MTS | 2 |
| 18 | F | 12.1 | L | 6.4 | NDD | 5 | 58 | F | 40.5 | R | 32.6 | SVM | 5 |
| 19 | M | 12.9 | R | 9.6 | FCD | 2 | 59 | M | 40.7 | R | 30.3 | LGT | 4 |
| 20 | F | 13.6 | R | 13.6 | HME | 8 | 60 | F | 41.2 | R | 2.2 | FCD | 4 |
| 21 | F | 14.4 | R | 14.4 | FCD | 24 | 61 | F | 41.6 | L | 11.1 | LGT | 2 |
| 22 | F | 15.1 | R | 10.1 | FCD | 14 | 62 | M | 43.1 | R | 37.0 | MTS | 2 |
| 23 | F | 15.2 | L | 14.6 | FCD | 13 | 63 | M | 43.5 | R | 19.0 | MTS | 5 |
| 24 | M | 15.2 | R | 6.9 | FCD | 10 | 64 | M | 43.6 | L | 39.6 | MTS | 5 |
| 25 | M | 15.5 | L | 9.5 | UNK | 16 | 65 | M | 43.8 | L | 8.8 | MTS | 5 |
| 26 | M | 17.8 | R | 16.8 | MTS | 5 | 66 | F | 44.9 | R | 22.2 | MTS | 9 |
| 27 | M | 19.1 | R | 2.2 | SVM | 1 | 67 | M | 46.3 | L | 31.3 | NA | 13 |
| 28 | M | 19.9 | L | 4.0 | SVM | 5 | 68 | F | 46.3 | R | 29.7 | SVM | 2 |
| 29 | M | 20.3 | R | 4.6 | LGT | 5 | 69 | M | 46.8 | R | 12.1 | SVM | 1 |
| 30 | F | 20.3 | R | 16.7 | SVM | 3 | 70 | F | 47.0 | R | 21.2 | LGT | 6 |
| 31 | M | 21.2 | L | 13.3 | LGT | 4 | 71 | F | 48.3 | R | 1.8 | SVM | 2 |
| 32 | M | 22.1 | R | 5.1 | NA | 6 | 72 | M | 49.2 | L | 41.1 | MTS | 5 |
| 33 | M | 24.8 | L | 6.2 | SVM | 3 | 73 | M | 49.2 | R | 33.0 | MTS | 1 |
| 34 | M | 24.8 | L | 16.0 | MTS | 4 | 74 | F | 51.0 | R | 45.0 | MTS | 2 |
| 35 | M | 25.0 | R | 10.0 | SVM | 7 | 75 | M | 51.4 | R | 48.0 | SVM | 1 |
| 36 | M | 25.4 | L | 1.9 | LGT | 12 | 76 | M | 52.7 | L | 34.7 | LGT | 8 |
| 37 | M | 26.5 | L | 8.0 | NDD | 9 | 77 | M | 52.8 | L | 47.0 | MTS | 3 |
| 38 | F | 27.4 | R | 13.4 | MTS | 5 | 78 | F | 52.9 | L | 42.4 | MTS | 3 |
| 39 | M | 27.7 | L | 3.6 | LGT | 3 | 79 | M | 54.2 | L | 9.0 | SVM | 3 |
| 40 | M | 29.2 | R | 6.6 | SVM | 4 | 80 | F | 54.4 | R | 29.4 | SVM | 1 |

**Table S2.** Dataset composition. Cell and subject count for every dataset used in this study. The first row for each species is the core dataset, the original temporal cortex recordings on which all developmental models are fitted. Rows marked Additional support further analyses indicated; all mouse data are from C57BL/6J animals except those of Oswald and Reyes, which are from Swiss Webster. Raw recordings under source data indicates all data was processed through same feature extraction and quality control pipeline as core dataset.

|  |  | Dataset | Sources (cells \| subjects) | Source data | Cells \| subjects | Age range | Presented | Species total |
| --- | --- | --- | --- | --- | --- | --- | --- | --- |
| Human |  | Temporal cortex | This study | Raw recordings | 584 \| 80 | 0.91–54.4 yr | Throughout | 1055 \| 178*<br>629 original |
|  | Additional | Infant extension (Frontal, Occipital, Temporal & Parietal cortex) | Barzó et al. (67 \| 7), this study (6 \| 2) | Raw recordings | 73 \| 9 | 0.08–1.29 yr | Fig. S9 |  |
|  |  | Frontal cortex | Barzó et al. (120 \| 27), this study (45 \| 11), Allen Cell Types Database (27 \| 6) | Raw recordings | 192 \| 44 | 0.10–51 yr | Fig. S17a, SHERPA |  |
|  |  | Patch-seq reference (Temporal cortex) | Berg et al. (242 \| 50) | Raw recordings | 242 \| 50 | 18–69 yr | Fig. 3j,k, Fig. S15 |  |
| Mouse |  | Temporal association cortex | This study | Raw recordings | 151 \| 42 | P0.86–P86 | Throughout | 462 \| 158<br>162 original |
|  | Additional | Frontal cortex | Kroon et al. (41 \| 18), this study (11 \| 5) | Raw recordings | 52 \| 23 | P0.8–P30 | SHERPA |  |
|  |  | Auditory cortex | Oswald and Reyes | Cell-level values | 199 \| 50 | P9–P29 | SHERPA |  |
|  |  | Visual cortex | Ciganok-Hückels et al. | Raw recordings | 60 \| 43 | P10–P64 | SHERPA |  |

Data sources: Barzó et al., eLife (2025); Allen Cell Types Database; Berg et al., Nature (2021); Kroon et al., Sci Rep (2019); Oswald & Reyes, J Neurophysiol (2008); Ciganok-Hückels et al., Cereb Cortex (2022). \*De-duplicated total, frontal cortex and infant extension dataset share samples.

**Table S3.**
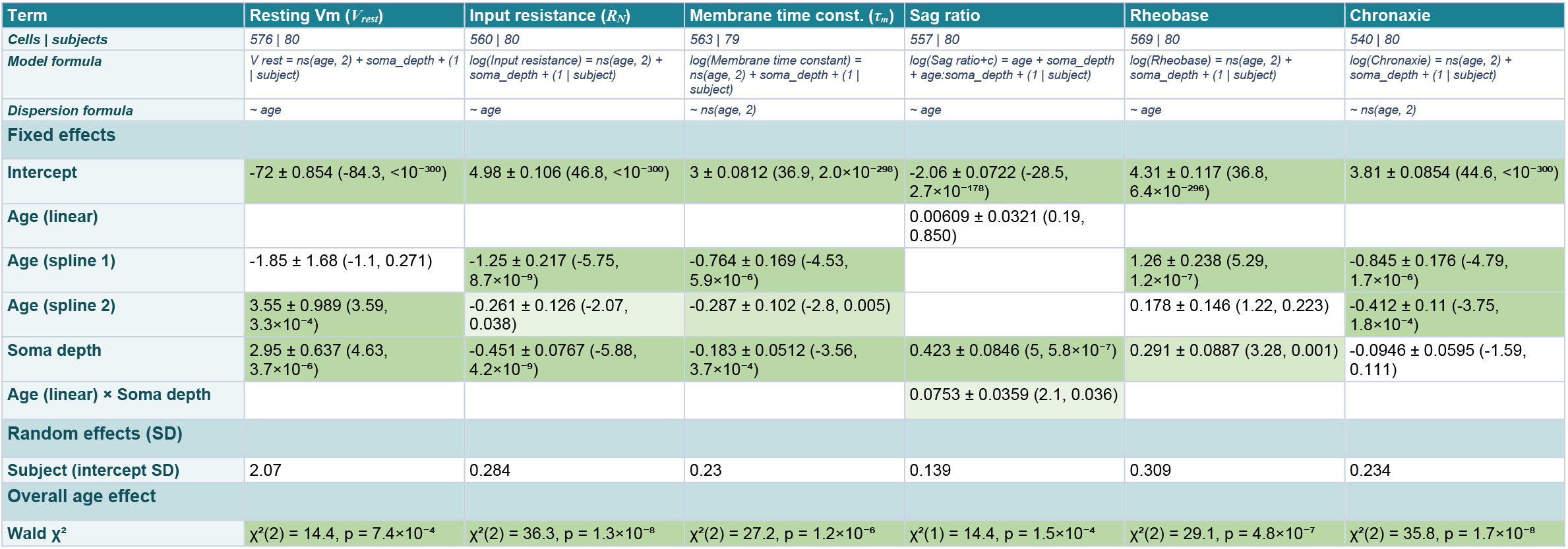
Human subthreshold properties and excitability. GAMM model specification and parameter estimates. Here and in following tables, parameter estimates presented as: estimate ± SE (z, p). Green shading marks significance in three levels (light p < 0.05; medium p < 0.01; intense p < 0.001). Blank cell means term not included in model.

**Table S4.**
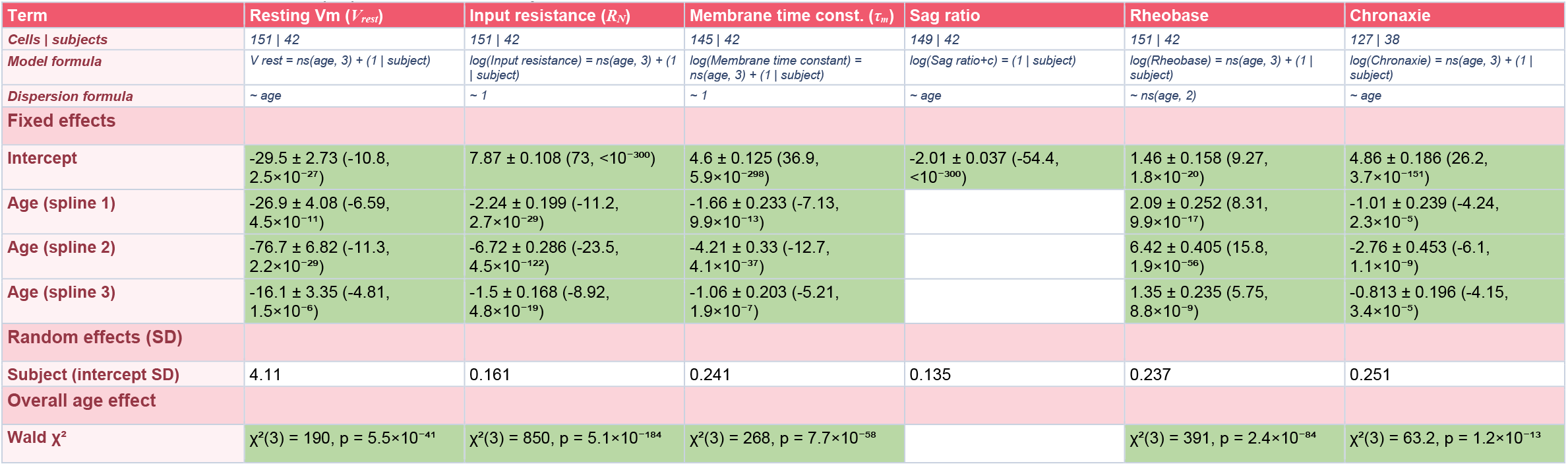
Mouse subthreshold properties and excitability. As Table S3, for mouse subthreshold properties and excitability.

**Table S5.** Human action potential waveform. Model specifications and parameter estimates for the action potential waveform features, presented as in Table S3. Because these features were measured on successive action potentials within each evoked spike train, the models include action potential number (AP1,AP10) as an additional fixed effect, and a random intercept for cell nested within subject.

| Term | AP threshold | AP amplitude | AP half-width | AP upstroke velocity | AP downstroke velocity |
| --- | --- | --- | --- | --- | --- |
| Cells \| subjects | 488 \| 68 | 488 \| 68 | 488 \| 68 | 488 \| 68 | 488 \| 68 |
| Model formula | $AP\ threshold = ns(age, 2) + AP\_nr + soma\_depth + (1 \vert subject/cell)$ | $AP\ amplitude = ns(age, 2) + AP\_nr + soma\_depth + (1 \vert subject/cell)$ | $log(AP\ half-width) = ns(age, 2) + AP\_nr + soma\_depth + (1 \vert subject/cell)$ | $log(AP\ upstroke\ velocity) = ns(age, 2) + AP\_nr + soma\_depth + (1 \vert subject/cell)$ | $AP\ downstroke\ velocity = AP\_nr + ns(age, 2) + soma\_depth + AP\_nr.ns(age, 2) + ns(age, 2).soma\_depth + (1 \vert subject/cell)$ |
| Dispersion formula | ~ age + AP_nr | ~ ns(age, 2) + AP_nr | ~ ns(age, 2) + AP_nr | ~ ns(age, 2) + AP_nr | ~ age |
| Fixed effects |  |  |  |  |  |
| Intercept | -37.2 ± 0.997 (-37.3, <10<sup>-300</sup>) | 78.8 ± 1.83 (43.2, <10<sup>-300</sup>) | 0.149 ± 0.0519 (2.87, 0.004) | 5.5 ± 0.0791 (69.5, <10<sup>-300</sup>) | -68.2 ± 4.81 (-14.2, 1.3×10<sup>-45</sup>) |
| Age (spline 1) | -6.4 ± 2.1 (-3.05, 0.002) | 15.7 ± 4.09 (3.83, 1.3×10<sup>-4</sup>) | -0.356 ± 0.116 (-3.06, 0.002) | 0.288 ± 0.177 (1.63, 0.104) | -28.8 ± 12.4 (-2.33, 0.020) |
| Age (spline 2) | -1.96 ± 1.36 (-1.44, 0.149) | 9.07 ± 2.54 (3.57, 3.6×10<sup>-4</sup>) | -0.132 ± 0.0724 (-1.83, 0.068) | 0.115 ± 0.11 (1.04, 0.297) | 6.79 ± 9.71 (0.7, 0.484) |
| Soma depth | -1.54 ± 0.746 (-2.07, 0.039) | 5.7 ± 1 (5.68, 1.3×10<sup>-8</sup>) | -0.0151 ± 0.0284 (-0.533, 0.594) | 0.352 ± 0.0427 (8.25, 1.5×10<sup>-16</sup>) | -2.87 ± 4.76 (-0.603, 0.547) |
| AP number (10) | 9.52 ± 0.389 (24.5, 4.2×10<sup>-132</sup>) | -16.5 ± 0.693 (-23.8, 9.7×10<sup>-125</sup>) | 0.521 ± 0.0177 (29.4, 1.5×10<sup>-189</sup>) | -0.64 ± 0.0263 (-24.3, 1.3×10<sup>-130</sup>) | 31.4 ± 1 (31.2, 6.3×10<sup>-214</sup>) |
| Age (spline 1) × AP number (10) | -8.43 ± 0.911 (-9.26, 2.1×10<sup>-20</sup>) | 15.5 ± 1.58 (9.85, 6.6×10<sup>-23</sup>) | -0.469 ± 0.0406 (-11.6, 6.9×10<sup>-31</sup>) | 0.65 ± 0.06 (10.8, 2.3×10<sup>-27</sup>) | -15.5 ± 2.6 (-5.95, 2.7×10<sup>-9</sup>) |
| Age (spline 1) × Soma depth |  |  |  |  | -13.3 ± 12.3 (-1.08, 0.281) |
| Age (spline 2) × AP number (10) | -3.88 ± 0.448 (-8.65, 5.1×10<sup>-18</sup>) | 7.71 ± 0.754 (10.2, 1.6×10<sup>-24</sup>) | -0.243 ± 0.0201 (-12.1, 1.6×10<sup>-33</sup>) | 0.352 ± 0.0284 (12.4, 2.8×10<sup>-35</sup>) | -13.1 ± 1.74 (-7.56, 4.0×10<sup>-14</sup>) |
| Age (spline 2) × Soma depth |  |  |  |  | -28 ± 9.24 (-3.04, 0.002) |
| Random effects (SD) |  |  |  |  |  |
| Subject (intercept SD) | 2.61 | 5.52 | 0.157 | 0.24 | 10.8 |
| Cell within subject (intercept SD) | 3.74 | 5.18 | 0.147 | 0.22 | 11.1 |
| Overall age effect |  |  |  |  |  |
| Wald $\chi^2$ | $\chi^2(2) = 11.8, p = 0.003$ | $\chi^2(2) = 28.5, p = 6.4 \times 10^{-7}$ | $\chi^2(2) = 13.2, p = 0.001$ | $\chi^2(2) = 3.87, p = 0.144$ | $\chi^2(2) = 25.4, p = 3.1 \times 10^{-6}$ |

**Table S6.**
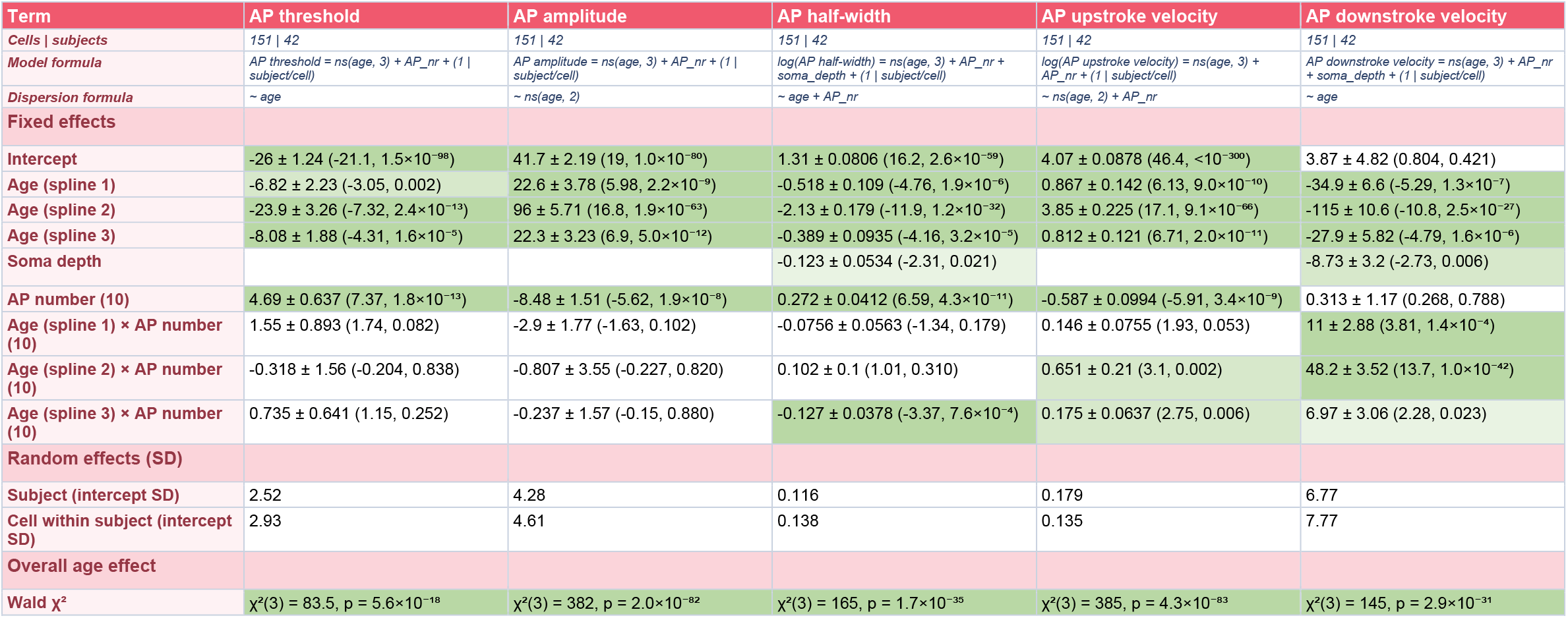
Mouse action potential waveform. As Table S5, for mouse action potential waveform.

**Table S7.**
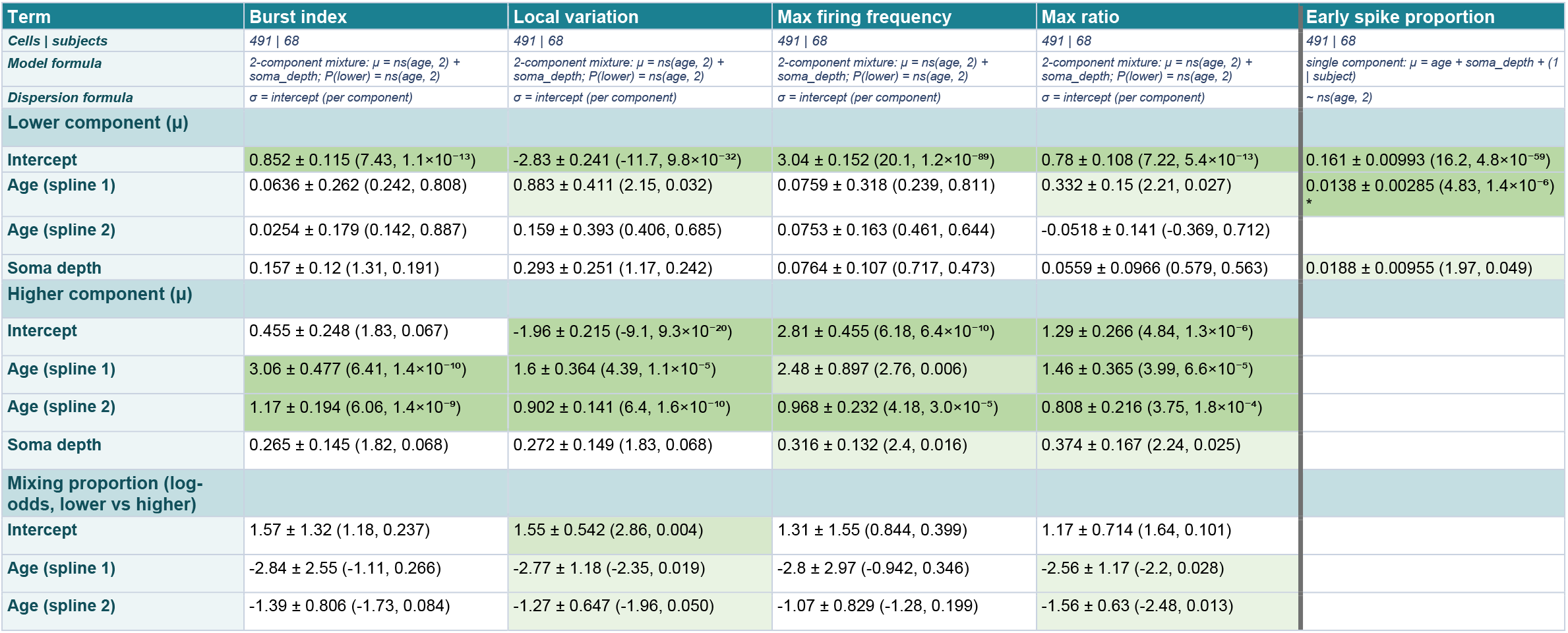
Human firing properties. Two-component GAMLSS mixture model specification for human firing properties. Early spike proportion is exception and fit with regular GAMM.

**Table S8.** Mouse firing properties. GAMM model specification for mouse firing properties.

| Term | Burst index | Local variation | Max firing frequency | Max ratio | Early spike proportion |
| --- | --- | --- | --- | --- | --- |
| Cells \| subjects | 122 \| 42 | 122 \| 42 | 122 \| 42 | 122 \| 42 | 122 \| 42 |
| Model formula | Burst index = ns(age, 2) + (1 \| subject) | Local variation = age + (1 \| subject) | Max firing frequency = ns(age, 3) + (1 \| subject) | Max ratio = age + (1 \| subject) | Early spike proportion = age + (1 \| subject) |
| Dispersion formula | ~ ns(age, 2) | ~ 1 | ~ ns(age, 2) | ~ 1 | ~ ns(age, 2) |
| Fixed effects |  |  |  |  |  |
| Intercept | 1.67 ± 0.17 (9.82, 9.4×10<sup>-23</sup>) | 0.0351 ± 2.3×10<sup>-4</sup> (155, <10<sup>-300</sup>) | 18.7 ± 1.66 (11.3, 1.7×10<sup>-29</sup>) | 1.92 ± 0.0416 (46.2, <10<sup>-300</sup>) | 0.12 ± 0.0153 (7.84, 4.5×10<sup>-15</sup>) |
| Age (linear) |  | -0.00363 ± 8.4×10<sup>-4</sup> (-4.32, 1.6×10<sup>-5</sup>) |  | -0.0753 ± 0.0232 (-3.25, 0.001) | 3.0×10<sup>-4</sup> ± 0.00381 (0.0792, 0.937) |
| Age (spline 1) | -0.625 ± 0.365 (-1.71, 0.087) |  | -1.93 ± 1.84 (-1.05, 0.294) |  |  |
| Age (spline 2) | 0.115 ± 0.201 (0.57, 0.569) |  | -17.4 ± 3.69 (-4.7, 2.6×10<sup>-6</sup>) |  |  |
| Age (spline 3) |  |  | -4.33 ± 1.59 (-2.72, 0.007) |  |  |
| Random effects (SD) |  |  |  |  |  |
| Subject (intercept SD) | 0.156 | 0.0168 | 1.62 | 0.291 | 0.0296 |

**Table S9.**
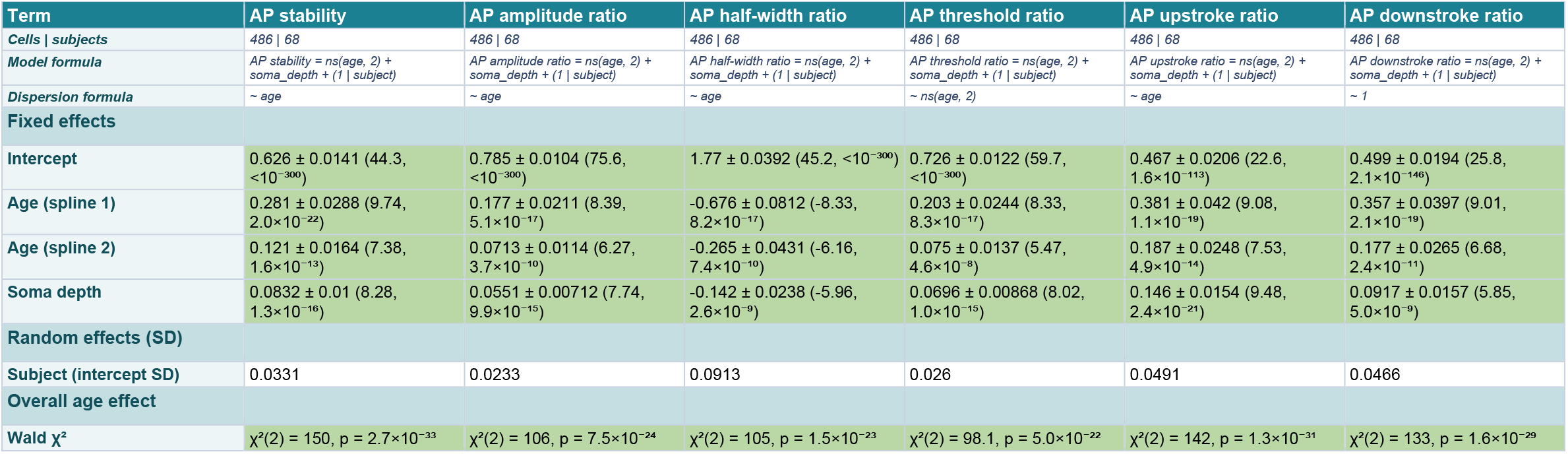
Human action potential waveform ratios and stability. GAMM model specification for human action potential waveform stability metrics

**Table S10.**
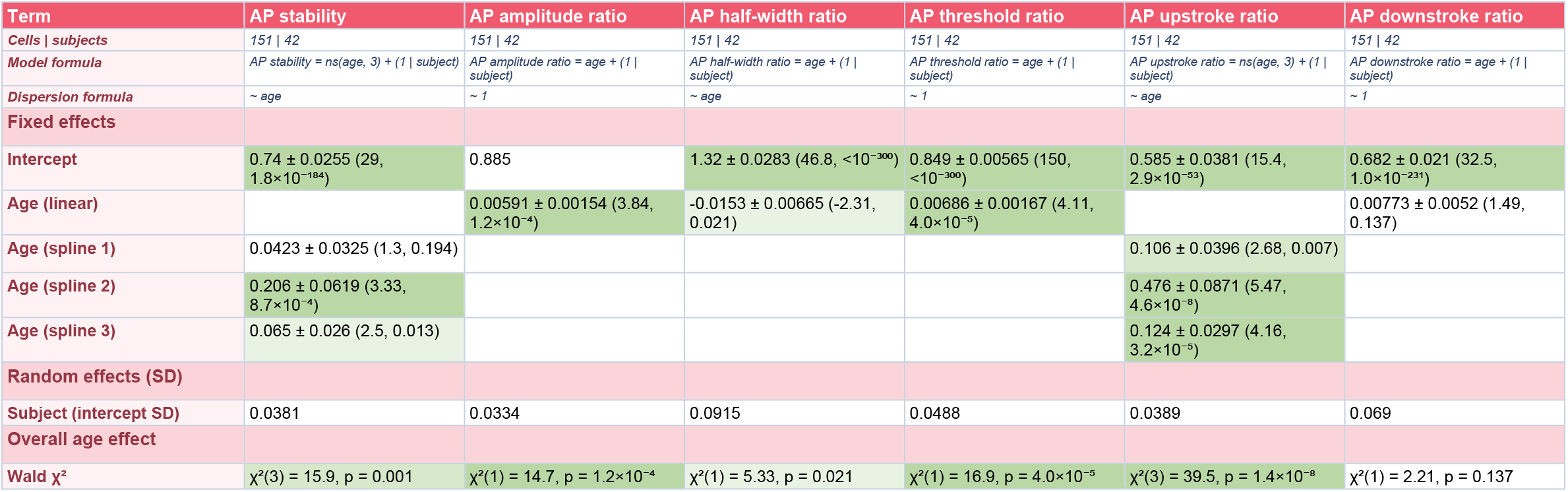
Mouse action potential waveform ratios and stability. As Table S9, for mouse action potential waveform stability metrics

**Table S11.** Leave-one-out cross-validation. Predictive accuracy assessed by leave-one-out cross-validation, in which each cell’s value was predicted from a model refitted with that cell held out, given its age and soma depth. R² and RMSE computed on fitted scale, median reports median of 12 features, AP features at AP 1.

| Feature | Scale | Human |  |  | Mouse |  |  |
| --- | --- | --- | --- | --- | --- | --- | --- |
|  |  | R<sup>2</sup> | RMSE | Cells \| donors | R<sup>2</sup> | RMSE | Cells \| animals |
| Input resistance ($R_{\text{in}}$) | log | 0.235 | 0.558 | 560 \| 80 | 0.887 | 0.438 | 151 \| 42 |
| Membrane time constant ($\tau_m$) | log | 0.129 | 0.39 | 563 \| 79 | 0.718 | 0.488 | 145 \| 42 |
| Sag ratio | original | 0.276 | 0.0858 | 557 \| 80 | 0.298 | 0.0591 | 149 \| 42 |
| Resting Vm ($V_{\text{rest}}$) | original | 0.022 | 5.68 | 576 \| 80 | 0.666 | 10.3 | 151 \| 42 |
| Rheobase | log | 0.182 | 0.609 | 569 \| 80 | 0.754 | 0.67 | 151 \| 42 |
| Chronaxie | fitted | 0.199 | 0.425 | 540 \| 80 | 0.277 | 0.555 | 127 \| 38 |
| AP amplitude (#1) | original | 0.242 | 7.52 | 488 \| 68 | 0.849 | 7.01 | 151 \| 42 |
| AP amplitude (#10) | original | 0.376 | 10.7 | 488 \| 68 | 0.756 | 9 | 151 \| 42 |
| AP half-width (#1) | log | 0.021 | 0.261 | 488 \| 68 | 0.818 | 0.195 | 151 \| 42 |
| AP half-width (#10) | log | 0.209 | 0.333 | 488 \| 68 | 0.719 | 0.254 | 151 \| 42 |
| AP upstroke velocity (#1) | log | 0.095 | 0.359 | 488 \| 68 | 0.824 | 0.299 | 151 \| 42 |
| AP upstroke velocity (#10) | log | 0.276 | 0.52 | 488 \| 68 | 0.751 | 0.422 | 151 \| 42 |
| AP downstroke velocity (#1) | original | 0.206 | 18.6 | 488 \| 68 | 0.829 | 11.1 | 151 \| 42 |
| AP downstroke velocity (#10) | original | 0.45 | 16.5 | 488 \| 68 | 0.622 | 12.7 | 151 \| 42 |
| AP threshold (#1) | original | 0.058 | 4.81 | 488 \| 68 | 0.609 | 3.67 | 151 \| 42 |
| AP threshold (#10) | original | 0.221 | 6.31 | 488 \| 68 | 0.402 | 5.15 | 151 \| 42 |
| AP stability index | original | 0.484 | 0.0813 | 486 \| 68 | 0.064 | 0.089 | 151 \| 42 |
| Median |  | 0.19 |  |  | 0.736 |  |  |

**Table S12.**
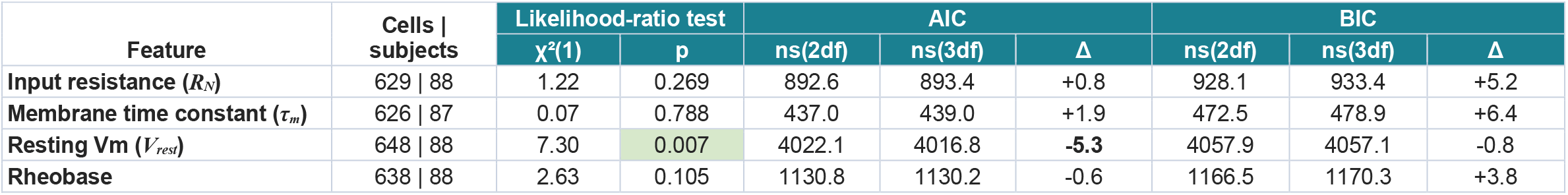
Model flexibility with extended infant data. Comparison of GAMMs fitted with 2 versus 3 degrees of freedom splines on age, with dataset extended to include more and younger infant data of mixed cortical origin (Table S2). Models are otherwise identical and compared by likelihood-ratio test.

**Table S13.**
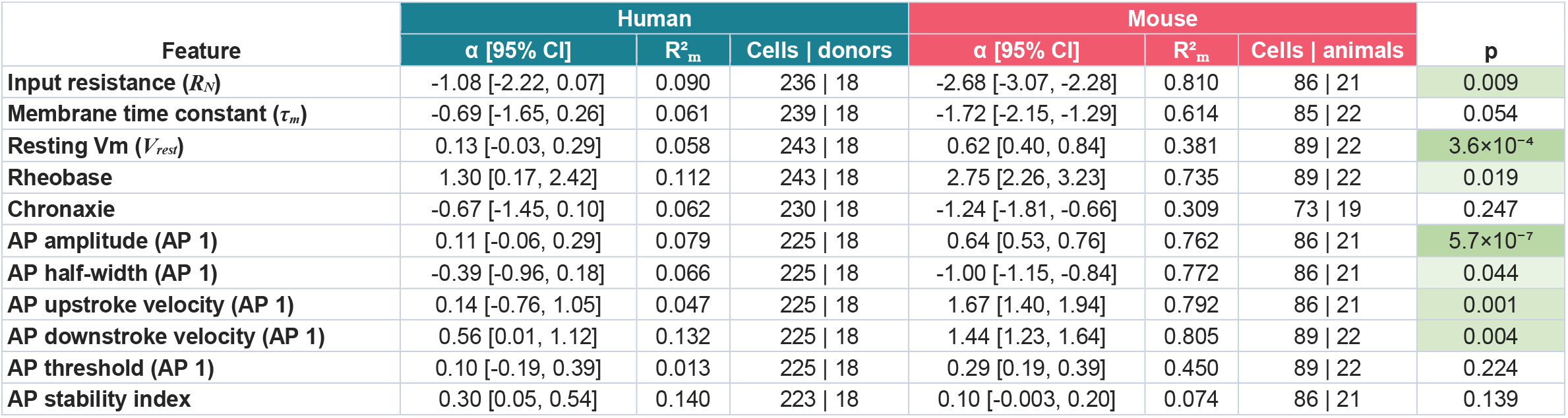
Allometric scaling of physiological features. Allometric scaling exponents (α) estimated by linear mixed models. The final column gives the p value for the species difference in α, shaded to level of significance as in Table S3.

| Feature | Human |  |  | Mouse |  |  | p |
| --- | --- | --- | --- | --- | --- | --- | --- |
| | $\alpha$ [95% CI] | $R^2_m$ | Cells \| donors | $\alpha$ [95% CI] | $R^2_m$ | Cells \| animals | |
| Input resistance ($R_N$) | -1.08 [-2.22, 0.07] | 0.090 | 236 \| 18 | -2.68 [-3.07, -2.28] | 0.810 | 86 \| 21 | 0.009 |
| Membrane time constant ($\tau_m$) | -0.69 [-1.65, 0.26] | 0.061 | 239 \| 18 | -1.72 [-2.15, -1.29] | 0.614 | 85 \| 22 | 0.054 |
| Resting Vm ($V_{rest}$) | 0.13 [-0.03, 0.29] | 0.058 | 243 \| 18 | 0.62 [0.40, 0.84] | 0.381 | 89 \| 22 | 3.6×10<sup>-4</sup> |
| Rheobase | 1.30 [0.17, 2.42] | 0.112 | 243 \| 18 | 2.75 [2.26, 3.23] | 0.735 | 89 \| 22 | 0.019 |
| Chronaxie | -0.67 [-1.45, 0.10] | 0.062 | 230 \| 18 | -1.24 [-1.81, -0.66] | 0.309 | 73 \| 19 | 0.247 |
| AP amplitude (AP 1) | 0.11 [-0.06, 0.29] | 0.079 | 225 \| 18 | 0.64 [0.53, 0.76] | 0.762 | 86 \| 21 | 5.7×10<sup>-7</sup> |
| AP half-width (AP 1) | -0.39 [-0.96, 0.18] | 0.066 | 225 \| 18 | -1.00 [-1.15, -0.84] | 0.772 | 86 \| 21 | 0.044 |
| AP upstroke velocity (AP 1) | 0.14 [-0.76, 1.05] | 0.047 | 225 \| 18 | 1.67 [1.40, 1.94] | 0.792 | 86 \| 21 | 0.001 |
| AP downstroke velocity (AP 1) | 0.56 [0.01, 1.12] | 0.132 | 225 \| 18 | 1.44 [1.23, 1.64] | 0.805 | 89 \| 22 | 0.004 |
| AP threshold (AP 1) | 0.10 [-0.19, 0.39] | 0.013 | 225 \| 18 | 0.29 [0.19, 0.39] | 0.450 | 89 \| 22 | 0.224 |
| AP stability index | 0.30 [0.05, 0.54] | 0.140 | 223 \| 18 | 0.10 [-0.003, 0.20] | 0.074 | 86 \| 21 | 0.139 |

**Table S14.** Allometric scaling with extended infant data. Linear mixed models as in Table S13, refitted for the features in the infant extension analysis. Each cell gives α, its 95% confidence interval in square brackets, and in parentheses the p value for the difference from the mouse exponent for that feature (Table S13), shaded to level of significance as in Table S3.

| Feature | Human, main | Human, with extended infant data |
| --- | --- | --- |
| Input resistance ($R_N$) | -1.08 [-2.22, 0.07] (0.009) | -0.79 [-1.30, -0.27] (7.5×10<sup>-9</sup>) |
| Membrane time constant ($\tau_m$) | -0.69 [-1.65, 0.26] (0.054) | -0.30 [-0.77, 0.17] (1.1×10<sup>-5</sup>) |
| Resting Vm ($V_{rest}$) | 0.13 [-0.03, 0.29] (3.6×10<sup>-4</sup>) | 0.18 [0.11, 0.25] (1.6×10<sup>-4</sup>) |
| Rheobase | 1.30 [0.17, 2.42] (0.019) | 1.15 [0.54, 1.77] (5.6×10<sup>-5</sup>) |

**Table S15.** Epilepsy etiology and duration effects. Likelihood-ratio tests comparing, for each feature, the final human GAMM (Tables S3, S5, S7 and S9) with the same model extended by (1) etiology group as a categorical fixed effect (Developmental: FCD, HME, NDD; Hippocampal sclerosis: MTS; Structural: SVM, LGT; Other/Unknown: non-lesional or unavailable etiology; abbreviations as in Table S1), (2) age-adjusted epilepsy duration residuals as a continuous predictor, and (3) the duration residuals together with their interaction with age. For each comparison the table gives the likelihood-ratio χ², its p value and the change in BIC of the extended model relative to the base model (negative values favour the extended model). Shading marks significance as in Table S3; n.c., the extended model did not converge.

| Feature | Cells \| subjects | Etiology group |  |  | Epilepsy duration |  |  | Duration × age |  |  |
| --- | --- | --- | --- | --- | --- | --- | --- | --- | --- | --- |
| | | $\chi^2$ | P | ΔBIC | $\chi^2$ | P | ΔBIC | $\chi^2$ | P | ΔBIC |
| Resting Vm ($V_{rest}$) | 576 \| 80 | 2.8 | 0.424 | +16.3 | 0.1 | 0.797 | +6.3 | 0.7 | 0.719 | +12.1 |
| Input resistance ($R_N$) | 560 \| 80 | 6.7 | 0.081 | +12.2 | 0.0 | 0.948 | +6.3 | 0.1 | 0.974 | +12.6 |
| Membrane time constant ($\tau_m$) | 563 \| 79 | 4.6 | 0.202 | +14.4 | n.c. | | | 1.2 | 0.544 | +11.4 |
| Sag ratio<sup>1</sup> | 557 \| 80 | 13.2 | 0.004 | +5.7 | 0.6 | 0.455 | +5.8 | 1.8 | 0.409 | +10.9 |
| Rheobase | 569 \| 80 | 4.6 | 0.202 | +14.4 | 1.1 | 0.289 | +5.2 | 1.2 | 0.555 | +11.5 |
| Chronaxie | 540 \| 80 | 2.6 | 0.461 | +16.3 | 0.0 | 0.863 | +6.3 | 0.4 | 0.819 | +12.2 |
| AP threshold | 488 \| 68 | 5.4 | 0.142 | +15.2 | 1.7 | 0.188 | +5.2 | 2.6 | 0.273 | +11.2 |
| AP amplitude | 488 \| 68 | 6.6 | 0.086 | +14.1 | n.c. |  |  | 2.1 | 0.352 | +11.7 |
| AP half-width | 488 \| 68 | 1.2 | 0.749 | +19.4 | 0.4 | 0.527 | +6.5 | 2.6 | 0.277 | +11.2 |
| AP upstroke velocity | 488 \| 68 | 3.2 | 0.359 | +17.4 | 1.0 | 0.309 | +5.9 | n.c. |  |  |
| AP downstroke velocity | 488 \| 68 | 3.2 | 0.358 | +17.4 | 0.3 | 0.613 | +6.6 | 1.9 | 0.387 | +11.9 |
| AP stability index<sup>2</sup> | 486 \| 68 | 11.1 | 0.011 | +7.5 | 0.2 | 0.630 | +6.0 | 0.6 | 0.757 | +11.8 |

<sup>1</sup>Developmental vs Structural, pairwise contrast\*: 0.210 ± 0.066 (z = 3.20, P = 0.008); Hippocampal sclerosis vs Structural: 0.175 ± 0.064 (z = 2.73, P = 0.032)
<sup>2</sup>Hippocampal sclerosis vs Developmental, pairwise contrast: 0.054 ± 0.018 (z = 2.95, P = 0.017); Other/Unknown vs Developmental: 0.047 ± 0.018 (z = 2.65, P = 0.040)
\*Tukey-adjusted pairwise contrasts between etiology groups (estimate ± SE on the model scale, with z and P)

